# Parvalbumin interneuron maturation reorganizes cortical population activity structure during development

**DOI:** 10.64898/2026.10.22.761282

**Authors:** Michelle W. Wu, Matteo Mariani, Anand Suresh, Nazim Kourdougli, Grace Jiang, Akash J. Kothari, Tanya S. Jain, Mario Dipoppa, Carlos Portera-Cailliau

## Abstract

During brain development, spontaneous cortical activity undergoes a dramatic transformation from highly synchronous network events to less correlated and sparse neuronal firing that is thought to underlie more flexible neural coding. However, how the structure of population activity changes across this transition, what circuit mechanisms drive this reorganization, and the implications for information encoding have not been fully established. Using longitudinal in vivo calcium imaging in early postnatal mice, we demonstrate that cortical network desynchronization during the second postnatal week is accompanied by a redistribution of fractional variance from dominant, low-dimensional modes into many weaker dimensions, and a growing influence of GABAergic interneurons (INs) on the network. All-optical circuit interrogation revealed that parvalbumin (PV) INs, but not somatostatin (SST) INs, are sufficient to desynchronize the network and shift the distribution of population variance. Through targeted ablation of a fraction of PV INs, we show dimensionality is highly correlated with stimulus decoding accuracy. Furthermore, PV INs undergo a sharp increase in transcriptional maturation between postnatal day (P) 8 and 11, during which they preferentially enrich for synaptic genes. Lastly, a novel, biologically plausible, multi-cell-type computational model reproduced the developmental desynchronization and rise in dimensionality. Strikingly, without being constrained by perturbation data, the model accurately predicted the effects of each IN class on the network that we observed empirically.

## INTRODUCTION

Spontaneous network activity in the cerebral cortex undergoes a dramatic transformation during development from an initial pattern that is dominated by intermittent, highly synchronous bursts of neuronal firing to a sparse and continuous pattern that is typical of the mature brain^1–4^. In rodents, this transition occurs during the second postnatal week and is characterized by reduced pairwise correlations between neurons, an increased frequency of population events, and a decreased neuronal participation in those events^5–9^. This developmental transformation of cortical network dynamics is thought to be of fundamental importance because of its implications for neural computation^4,5,10,11^. However, although the phenomenology of this network transition and the contributions of sensory input in generating synchronous bursts have been well characterized^12–15^, neither the circuit mechanisms that underlie this transition nor its consequences for neural population coding have been elucidated.

One influential framework for cortical computation proposes that primary sensory cortices implement sparse and distributed population codes^16–18^, which are predicted to support a high-dimensional representational space with large coding capacity^19^. In the adult cortex, neural population activity occupies a high-dimensional state space, with both sensory-evoked and spontaneous activity exhibiting variance distributed across many dimensions^20,21^. Although pairwise correlations decrease during development, it remains unclear how this transition gives rise to the high-dimensional population activity characteristic of the mature cortex. Recent work has shown that spontaneous cortical activity becomes higher-dimensional across the second postnatal week^22^, suggesting that this developmental transition involves a reorganization of the structure of population activity. How this reorganization emerges mechanistically, and its consequences for the cortex’s coding capacity, remain unknown.

A leading candidate mechanism for this developmental reorganization is the maturation of cortical inhibitory circuits. In particular, GABAergic inhibitory interneurons (INs) born in the medial ganglionic eminence (MGE) undergo a prolonged maturation process that extends postnatally^23–25^, coinciding with the developmental window during which cortical population activity decorrelates. The MGE gives rise to two major classes of IN, parvalbumin (PV) and somatostatin (SST), which express an overlapping sequence of transcription factors throughout their embryonic and early postnatal development in mice^26–28^. Despite this shared lineage, PV and SST INs target different compartments of excitatory (Exc) neurons to differentially shape cortical network activity^29^. Recent studies have begun to uncover how manipulating inhibitory activity affects network synchrony, such that broad optogenetic activation of GABAergic INs decorrelates spontaneous cortical activity^30^ while chemogenetic activation of SST INs increases spontaneous correlations^31^. However, whether specific IN subtypes drive the developmental expansion of population dimensionality has never been explored. Here, we investigate how the structure of population activity reorganizes across development and identify the circuit and cellular mechanisms that drive the emergence of high-dimensional cortical activity.

Because interneuron (IN) subtypes act within recurrent circuitry, a computational model is crucial to establish how each subtype contributes to developmental desynchronization. However, no existing mechanistic model combines all the features needed to address this problem: recurrent connectivity, single-neuron resolution, IN-subtype specificity, and slow population fluctuations at the correlation levels measured over the second postnatal week. In fact, several existing models of developmental changes in that time period are feedforward and thus lack recurrent connectivity^32–34^ or single-cell resolution^35,36^. Meanwhile, population models with single-neuron resolution did not differentiate between IN subtypes^30,37^ or, in their mechanistic versions, did not consider a sufficiently long span of developmental age and fluctuations generated intrinsically by the network^38^. Existing recurrent multi-cell-type models of intrinsic neural synchrony with single-neuron resolution^39,40^ focus on the adult mouse brain and consistently show faster fluctuations than those observed in development. Conversely, models with emergent slow fluctuations in neural activity either lack the single-neuron resolution needed to study dimensionality of neural activity^41^ or they require synchronous events to be driven exogenously^42^. Finally, single-neuron resolution models capable of generating slow-rate fluctuations exhibit average neuronal correlations that are much weaker than those observed experimentally^43^. Together, these limitations motivate us to develop a recurrent, multi-cell-type circuit framework that can simultaneously capture the slow dynamics of SNEs, the structure of population activity, and the experimentally observed levels of correlation, enabling us to investigate which circuit mechanisms drive the reorganization of cortical dynamics.

By leveraging longitudinal in vivo two-photon calcium imaging of excitatory and MGE INs, we found that population activity increases its dimensionality during the second postnatal week through a redistribution of fractional variance (the proportion of population activity patterns explained by each dimension) from a few dominant dimensions to many weaker ones. Optogenetic activation of PV but not SST INs at P10 partially recapitulated the restructuring of population activity. Through single-nucleus RNA-sequencing analyses, we identified P8-P11 as a major transition period for the transcriptional maturation of MGE INs, during which PV and SST INs manifest unique gene-level implementations within shared pathways. We also link population dimensionality to stimulus encoding. Finally, we developed a multi-cell-type recurrent circuit model of early postnatal cortical network activity that recapitulates major experimentally observed features of developmental network reorganization, predicts qualitatively the effects of PV IN activation observed in the data, and identifies circuit mechanisms that can account for coordinated changes in network dynamics.

## RESULTS

### Population activity restructures during the second postnatal week

To record the activity of excitatory (Exc) and inhibitory (Inh) neurons in S1, we performed longitudinal in vivo two-photon calcium imaging (GCaMP8s, 15.1 Hz) from P9 to P14 in head-fixed Nkx2.1-Cre;Ai14 mice that express tdTom in MGE-derived INs^44^ (**Figures 1A** and **S1A**). At P9, network activity appeared as episodic bursts of activity involving large numbers of both Exc and Inh neurons, whereas at P14 activity was continuous and sparse (**Figure 1B**). Quantification of the properties of synchronous network events (SNEs), defined as periods in which a higher-than-chance proportion of neurons are simultaneously active, showed that their frequency significantly increased from P9 to P13 (**Figure 1C**), while their duration decreased (**Figure S1B**). During this period, the fraction of Exc neurons recruited per SNE was reduced by half, from ∼40% at P9 to ∼20% at P13 (**Figure 1D**). In contrast, Inh neurons were more globally and stably recruited across the second postnatal week, with 60-80% of the population participating in SNEs regardless of age (**Figure 1D**). Both the Exc population, despite its decline in recruitment, and the Inh population become more heterogeneous in their participation across SNEs at older ages, as reflected by an increasing coefficient of variation (**Figure S1C**), suggesting that the neurons being recruited are increasingly variable across SNEs.

**Figure 1.**
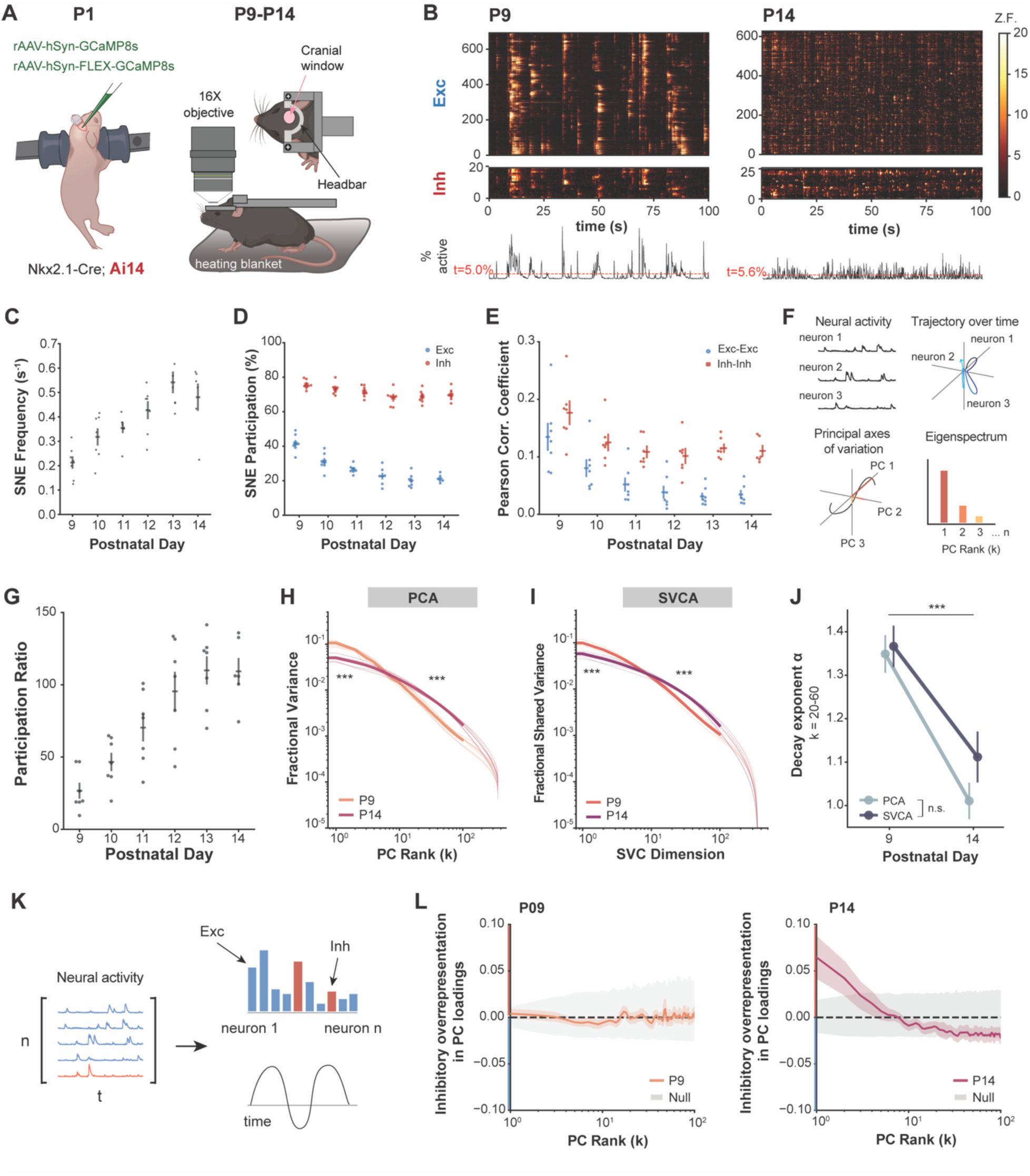
Developmental restructuring of cortical population activity during the second postnatal week. (A) Experimental design for longitudinal 2-photon calcium imaging. (B) Top: Example raster of spontaneous activity of Exc and Inh neurons, sorted with Rastermap^93^, at P9 and P14. Bottom, percentage of the total population that is active. Dashed red line indicates statistical threshold for SNEs. Modified Z-scored fluorescence used for heatmaps. (C) SNE frequency (events per second) increases across the second postnatal week. Linear mixed-effects model (LMM) shows significant effect of Day (Wald χ^2^ = 142.41, p = 5.50x10^-29^). N = 7 mice for P9-P13, n = 6 for P14, 500-900 neurons per FOV. (D) Proportion of Exc and Inh neurons participating in SNEs at different ages. Significant main effects of Day (Wald χ^2^ = 360.70, p = 8.71x10^-76^), Cell Type (Wald χ^2^ = 151.35, p = 8.81x10^-35^), and a significant Day x Cell Type interaction (Wald χ^2^ = 136.57, p = 9.60x10^-28^). Post hoc pairwise comparisons between Exc and Inh within each postnatal day were corrected for multiple comparisons and were significant for all days (adjusted p < 0.001). (E) Pearson correlation coefficient (PCC) of Exc-Exc and Inh-Inh neuron pairs. Significant main effects of Day (Wald χ^2^ = 191.80, p = 1.61x10^-39^), Cell Type (Wald χ^2^ = 13.56, p = 1.14x10^-3^), and a significant Day x Cell Type interaction (Wald χ^2^ = 46.95, p = 9.6x10^-7^). Post hoc pairwise comparisons between Exc and Inh within each postnatal day were significant for all days (adjusted p < 0.05). (F) Schematic of the relationship between spontaneous neural activity, trajectory geometry, dimensionality reduction, and the eigenspectrum in an example case with three neurons for visualization purposes (adapted from Pospisil and Pillow^94^). (G) Participation ratio of subsampled population (350 neurons, 200 repeats) increases over the second postnatal week. Significant effect of Day (Wald χ^2^ = 328.15, p = 8.82x10^-69^). (H) Fraction of variance explained by descending orders of principal components (PCs). For *h-j*, recordings were subsampled to 360 neurons (200 repeats); thin lines= individual mice, thick lines= mean. LMM reveals significant difference between P9 and P14 at early (PCs 1-5) and intermediate ranks (PCs 20-60). (Early: β = -0.592, p < 0.001; Intermediate: β = 0.719, p < 0.001). (I) Fraction of shared variance explained by descending orders of SVC dimensions. Significant difference between P9 and P14 at early and intermediate ranks (Early: β = -0.459, p < 0.001; Intermediate: β = 0.537, p < 0.001). (J) Decay exponent α estimated over the stable intermediate range of k = 20-60 for both PCA and SVCA eigenspectra. Significant main effect of Day on α (Wald χ^2^ = 32.35, p = 1.29x10^-8^) but no significant effect detected for Method (Wald χ^2^ = 0.066, p = 0.797). (K) Schematic illustrating the decomposition of population activity by PCA. Each principal component is characterized by a loading vector describing the contribution of each neuron (Exc, blue; Inh, red) and a time course describing that PC’s activity over time. **l,** Inhibitory overrepresentation in PC loadings as a function of PC rank at P9 (left) and P14 (right), relative to a null distribution generated by randomly permuting cell type labels (gray). Color shaded regions indicate mean ± SEM. Dashed line indicates 0 (proportional representation).

Consistent with previous work, we observed a decline in Exc-Exc pairwise correlations between P9 and P13^5,22^. This trajectory was significantly different than that of Inh-Inh pairwise correlations, which remained elevated after P10 (**Figure 1E**). Analyzing pairwise correlations as a function of distance between neurons revealed that the distance-dependence of Exc-Exc correlations decays more than that of the Inh-Inh correlations (**Figure S1D**), in accordance with previous findings^5,45^. Furthermore, Exc-Exc and Inh-Inh correlations were significantly greater than chance at both P9 and P14, although for Exc-Exc pairs the difference was much smaller at P14 than for Inh-Inh pairs (**Figure S1E**).

To characterize how spontaneous population activity is organized during development, we applied principal component analysis (PCA), which decomposes population activity into orthogonal modes (principal components, PC) that can be ordered by the fraction of variance they explain (fractional variance; eigenvalues normalized by total variance, i.e., the sum of all eigenvalues) (**Figure 1F**). From this decomposition we derived the participation ratio (PR), a summary statistic that estimates the effective dimensionality of a population by quantifying how evenly fractional variance is distributed across PCs^46^. The previously described age-dependent increase in SNE frequency and decrease in Exc-Exc correlations were accompanied by a dramatic increase in PR for the excitatory population (**Figure 1G**), indicating a developmental shift toward higher-dimensional activity. To examine this shift across the full dimensionality of the data, rather than the single value captured by PR, we next inspected the eigenspectrum, the normalized eigenvalues ranked in descending order, which describes how fractional variance is distributed across all dimensions of population activity (**Figure 1F**; see Methods). The increase in participation ratio (**Figure 1G**) indicates that the distribution of eigenvalues becomes less concentrated across development, but this could be consistent with different spectrum shapes.

To further investigate different possibilities, we first examined how the eigenspectrum of population activity changes between P9 and P14 using random subsamples of neurons (see Methods). At P9, fractional variance was concentrated in the leading dimensions, with a relatively steep decay across dimensions. In contrast, at P14, the fractional variance spectrum was less concentrated in the leading dimensions and more evenly distributed across dimensions, leading to a shallower eigenspectrum than at P9 (**Figure 1H**). Thus, fractional variance is redistributed from the leading dimensions into the later dimensions, producing a crossover in the eigenspectrum. To confirm that our subsample size was sufficient to capture spectral shape, we repeated this analysis across a range of subsample sizes (**Figure S2A**; see Methods). Although fractional variance showed more spread across subsampling levels at P14, spectra that were rescaled relative to the mean variance of the top 5 PCs (to enable shape comparison by aligning the spectra to the top components, which are most stable across subsampling levels) aligned well at early and intermediate ranks down to subsamples of just 180 neurons, indicating that subsamples of 180 neurons or larger reliably capture spectral shape. At both ages, spectra were unstable at the tail, reflecting proximity to the noise floor.

While PCA captures all variance, it does not distinguish between variance that is reproducible across time and variance that reflects transient or noisy neural activity. Therefore, we next applied a variation of shared variance component analysis (SVCA, modified from Stringer et al.^21^; see Methods) to estimate temporally reliable variance, defined as the variance shared across non-overlapping time windows (i.e., shared variance). Fractional shared variance was more evenly distributed across dimensions at P14 compared to at P9 (**Figure 1I**), reflecting the redistribution of fractional variance seen in the PCA eigenspectrum. In the adult cortex, the eigenspectrum and shared variance spectrum follow an approximate power law^20,21^. During the second postnatal week, the spectra are not linear on a log-log scale, suggesting that this organizational principle of mature population activity has not yet emerged. However, to quantify the difference in spectral shape between ages, we estimated the decay exponent (α) over an intermediate range of dimensions. The decay exponent was significantly smaller at P14 than at P9 for both PCA and SVCA (**Figure 1J**), indicating a shallower spectral decay and a progression towards a more distributed spectrum. Additionally, we found that the reliable variance fraction, defined as the ratio of SVCA to PCA fractional variance at matched ranks, increased broadly across all dimensions at P14 compared to P9 (**Figure S2B**), indicating that a larger fraction of overall population activity is temporally reliable across the spectrum. Thus, the circuit at P14 reorganizes how fractional shared variance is distributed across dimensions, resulting in a population that explores more dimensions and does so more consistently. Overall, these findings demonstrate that cortical network activity undergoes a profound transformation over just a few days during the second postnatal week, achieving a higher dimensionality consistent with the more distributed population geometry reported in the adult cortex^21^.

Whereas the eigenspectrum describes how much variance each PC explains, the PC loadings describe which neuron types, Exc or Inh, contribute (**Figure 1K**). Our recordings in Nkx2.1-Cre;Ai14 mice allowed us to examine the contributions of Exc and Inh neurons to each dimension. At P9, Exc and Inh neurons contributed to each PC in proportion to their numeric representation, consistent with the population variance having no cell type-specific organization (**Figure 1L**). In contrast, at P14, Inh neurons were overrepresented in the leading components. Both z-scored and mean-centered (**Figure S2C**; see Methods) analyses concur, suggesting that the dominant modes of population activity become preferentially shaped by inhibitory circuits across development regardless of variance in activity. This aligns with the timing of when MGE INs are integrated into cortical circuits during this postnatal period^3,23,27^. However, even though inhibition becomes more dominant in the leading PCs, this does not establish whether the redistribution of fractional variance is causally driven by inhibition, or which inhibitory subtype is responsible.

### PV interneuron activation decorrelates population activity and shifts fractional variance toward higher-dimensional modes

Previous studies had hinted at a role for GABAergic INs in mediating the desynchronization of network activity^30,31,47,48^. However, none of these prior studies examined the impact of such manipulations on population structure and dimensionality. To address this important question, we used acute cell type-specific optogenetic perturbations to determine whether transient activation of either of the MGE IN classes – PV or SST – is sufficient to reproduce the redistribution of variance observed between P9 and P14. To selectively target PV INs, we generated Nkx2.1-Cre;Sst-FlpO mice and injected rAAV8-nEF1-Con/Foff-ChRmine-oScarlet into S1 at P1 (see Methods, adapted from Kourdougli et al.^47^). We then acutely activated the excitatory opsin ChRmine in vivo by scanning a 1,040 nm laser across the field of view (6 trials, 10s on / 20s off) while simultaneously recording Exc and PV neurons at P10, P12, and P14 (**Figures 2A** and **2B**). ChRmine expression was already robust at P10, as indicated by oScarlet expression (**Figure S3A**). Opsin-expressing putative PV INs were identified by their time-locked responses to optogenetic stimulation and were excluded from subsequent population analyses of Exc neuron activity (**Figures S3B** and **S3C**).

**Figure 2.**
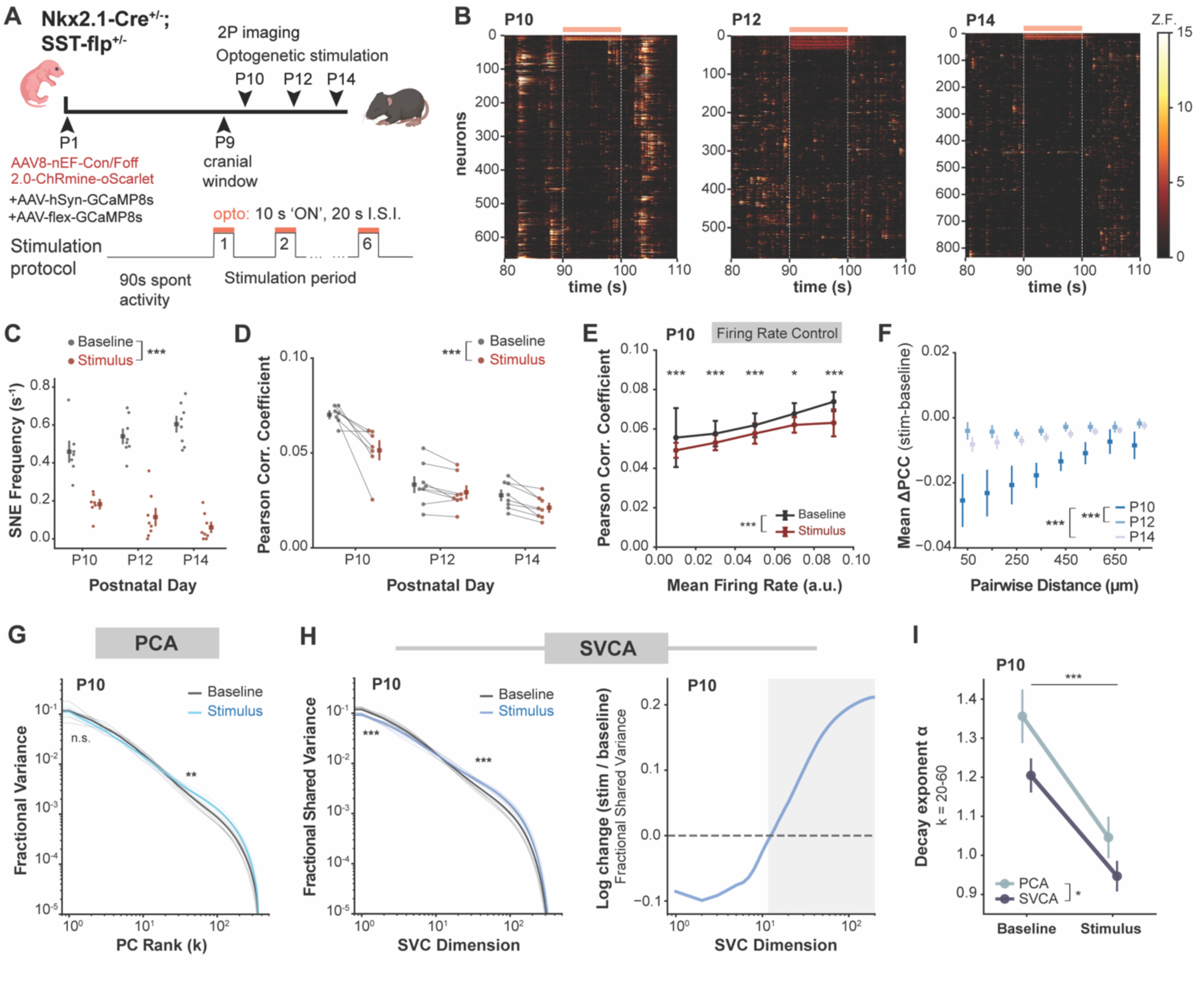
PV interneuron activation decorrelates network activity and shifts the distribution of variance. (A) Experimental design for optogenetic activation of PV interneurons (INs). (B) Example rasters (modified Z-scored fluorescence) showing 1 trial of activation at P10, P12, and P14. PV INs are sorted at the top, Exc neurons below. (C) SNE frequency during baseline vs. PV IN activation. LMM shows significant main effects of Day (Wald χ^2^ = 13.91, p = 9.54x10^-4^), Condition (Wald χ^2^ = 47.95, p = 4.37x10^-12^), and a significant Day x Condition interaction (Wald χ^2^ = 26.31, p = 1.94x10^-6^). Post-hoc contrasts showed significant effects of optogenetics at each day (adjusted p < 0.001), increasing over time (P10: *p* = 4.37×10^-12^; P12: *p* = 2.32×10^-35^; P14: *p* = 2.81×10^-42^). n=7-8 mice, 500-900 neurons per FOV. (D) Exc-Exc Pearson correlation coefficients (PCC) during baseline and during PV IN activation. Significant main effects of Day (Wald χ^2^ = 110.68, p = 9.24x10^-25^) and Condition (Wald χ^2^ = 15.70, p = 7.42x10^-5^). (E) Exc-Exc PCC binned by mean firing rate during baseline and PV IN activation periods. Points represent the mean across mice, and error bars indicate SEM across mice. Correlations were computed from an equal number of randomly sampled neuron pairs per recording (see Methods). Significant effects of Firing-Rate Bin (Wald χ^2^ = 47.71, p = 1.08x10^-9^), Condition (Wald χ^2^ = 124.58, p = 6.30x10^-29^), and a significant Firing-Rate Bin × Condition interaction (Wald χ^2^ = 57.90, p = 8.00x10^-12^) were observed. Post hoc LMMs were fit separately within each firing-rate bin to compare conditions (Baseline and Stimulus). The following firing-rate bins were significant (adjusted p < 0.05): 0.01, 0.03, 0.05, 0.07, 0.09. (F) Change in PCC as a function of distance between neuron pairs. Significant effect of Day (Wald χ^2^ = 32.52, p = 8.67x10^-8^). Relative to P10, change in PCC was significantly higher at P12 (β = 0.731, p < 0.001) and P14 (β = 0.545, p < 0.001). (G) Fraction of variance explained by descending ranks of PCs plotted on a log-log scale for baseline vs. PV IN activation (stimulus) at P10 (left). Recordings were subsampled to 360 neurons (200 repeats). Thin lines represent individual mice, thick lines represent the mean. Significant difference between baseline and PV IN activation at intermediate ranks (Early: β = - 0.122, p = 0.077; Intermediate: β = 0.177, p = 0.01). (H) Same as *g*, but fraction of shared variance explained by descending ranks of shared variance components (left). Significant difference between baseline and stimulation at early and intermediate ranks (Early: β = -0.210, p < 0.001; Intermediate: β = 0.272, p < 0.001). Ratio of PV IN activation and baseline fractional shared variance spectra (right). (I) Decay exponent α estimated over the stable intermediate range of k = 20-60 for both PCA and SVCA spectra. Significant main effect of Condition on α (Wald χ^2^ = 27.05, p = 1.98x10^-7^) and Method (Wald χ^2^ = 6.44, p = 0.011).

Acute activation of PV INs robustly suppressed network activity, significantly reducing SNE frequency during stimulation, an effect that became stronger across development (**Figure 2C**). Pairwise correlations were also significantly reduced during PV IN activation at all ages (**Figure 2D**). This reduction was largest at P10, even after controlling for the reduced firing rate of Exc neurons^49^ (**Figures 2E** and **S3D**). Additionally, at P10, but not at the older ages, the decrease in correlation was strongest for nearby neuron pairs, revealing a spatially localized effect of PV IN activation (**Figure 2F**). Differences in PV IN recruitment by optogenetic stimulation could not account for the effect on pairwise correlations, as activating even a small number of PV INs was sufficient to drive network desynchronization, and the percentage of stimulus responsive neurons did not predict the magnitude of the change in correlation (**Figure S3E**). Of note, the PV INs that we identified as being responsive to optogenetic activation likely represent only a subset of the true responsive population, since our assessment was restricted to cells that co-expressed ChRmine and GCaMP8s.

The observed effects on SNE frequency and pairwise correlations were also robust to the choice of protocol, as delivering optogenetic stimulation across 20 shorter trials (1s on / 3s off) produced similar reductions in SNE frequency and pairwise correlations (**Figures S4A-S4C**). Control experiments in animals expressing mCherry alone (no opsin) produced no change in SNE frequency or correlations during laser stimulation (**Figures S4D-S4F**), confirming that the observed effects are due to optogenetic activation of PV INs and not to optical artifacts.

Given that PV IN activation acutely desynchronizes cortical activity at P10, we next tested whether this manipulation is sufficient to reshape the structure of population activity similar to normal development. At P10, PV IN activation produced a significant increase in fractional variance in the intermediate-rank components, with no significant change in the leading components (**Figure 2G**). More strongly, PV IN activation significantly reduced the leading components and increased the intermediate-rank components of fractional shared variance (**Figure 2H**), in essence shifting its distribution in the direction of development. For both PCA and SVCA eigenspectra, the estimated decay exponent (α) over an intermediate range of dimensions was significantly smaller during PV IN activation (**Figure 2I**), indicating a more evenly distributed variance, just as we had observed during development (**Figure 1J**). Together, these results show that optogenetically engaging PV INs is sufficient to shift the distribution of fractional shared variance in the direction of the developmental transition during the second postnatal week.

The effect of PV IN activation on the distribution of fractional variance was greatest at P10, before the network desynchronizes, and was absent from the fractional variance eigenspectrum by P14, when cortical desynchronization is largely complete (**Figure S3F**). However, PV IN activation did produce a significant shift in fractional shared variance at P14, suggesting that PV IN activation at P14 modulates the reliability of activity within an already-redistributed framework, rather than further reorganizing the population structure itself.

### SST interneurons do not shift fractional variance

We next tested whether activation of SST INs produces similar effects on population activity, using Sst-FlpO mice and injecting a rAAV8-nEF1-Coff/Fon-ChRmine-oScarlet virus (**Figures 3A, 3B, S5A,** and **S5B**). Putative SST INs were identified by their reliable responses to optogenetic stimulation and were excluded from subsequent analyses. Similar to PV IN experiments, optogenetic activation of SST INs significantly reduced the frequency of SNEs across development, with larger effects at later ages (**Figure 3C**). However, SST IN activation did not consistently reduce pairwise correlations at any age (**Figures 3D, S5C,** and **S5D**). At P10, there was a modest increase in pairwise correlations at higher firing rates and for nearby neurons (**Figures 3E** and **3F**). Therefore, while SST IN activation suppressed network activity, it failed to desynchronize the population, underscoring the unique role of PV INs in shaping network synchrony during development.

**Figure 3.**
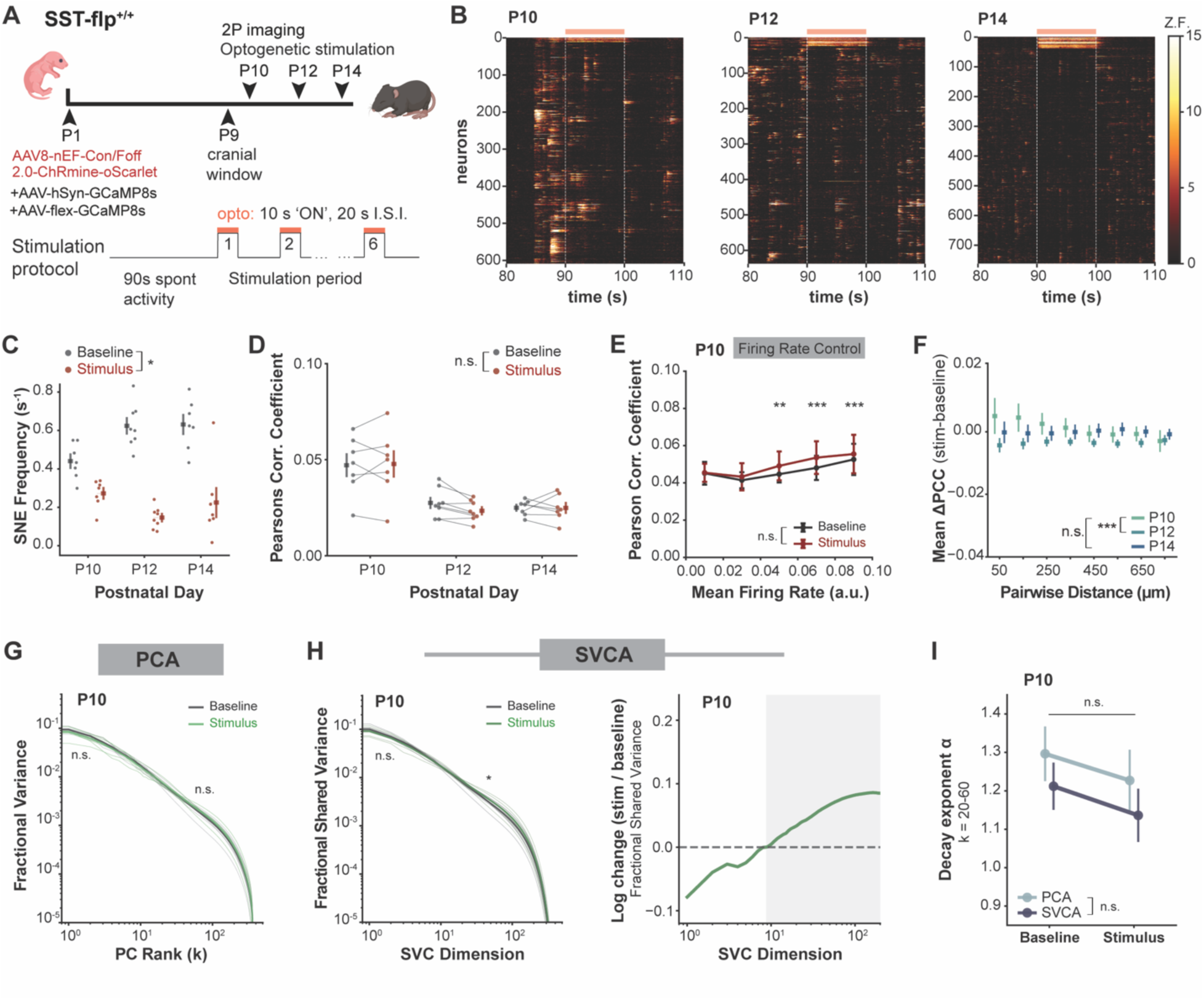
SST interneurons do not restructure population activity. (A) Experimental design for optogenetic activation of SST INs. (B) Example rasters showing 1 trial of activation at P10, P12, and P14. SST INs are sorted at the top, Exc neurons below. (C) SNE frequency during baseline vs. SST IN activation. LMM shows significant main effects of Day (Wald χ^2^ = 8.82, p = 0.012), Condition (Wald χ^2^ = 4.78, p = 0.029), and a significant Day x Condition interaction (Wald χ^2^ = 17.49, p = 1.59x10^-4^). Post hoc contrasts showed significant effects of optogenetics at each day (adjusted p < 0.05), increasing over time (P10: *p* = 0.029, P12: *p* = 3.43×10⁻^17^; P14: *p* = 4.10×10⁻^12^). n=7-8 mice, 500-900 neurons per FOV. (D) Exc-Exc PCC during baseline and during SST IN activation. Significant main effects of Day (Wald χ^2^ = 40.71, p = 1.45x10^-9^), but not Condition (Wald χ^2^ = 0.184, p = 0.668). (E) Exc-Exc PCC binned by mean firing rate during baseline and stimulus periods. Points represent the mean across mice, and error bars indicate SEM across mice. Correlations were computed from an equal number of randomly sampled neuron pairs per recording (see Methods). Significant effects of Firing-Rate Bin (Wald χ^2^ = 35.41, p = 3.83 x10^-7^) but not Condition (Wald χ^2^ = 1.66, p = 0.20), and a significant Firing-Rate Bin × Period interaction (Wald χ^2^ = 24.88, p = 5.33 x10^-5^) were observed. Post hoc LMMs were fit separately within each firing-rate bin to compare Baseline and Stimulus. The following firing-rate bins were significant (adjusted p < 0.05): 0.05, 0.07, 0.09. (F) Change in PCC as a function of distance between neuron pairs. Significant effect of Day (Wald χ^2^ = 12.42, p = 0.002). Relative to P10, change in PCC was significantly lower at P12 (β = -0.482, p < 0.001) but not P14 (β = -0.225, p = 0.121). (G) Fraction of variance explained by descending ranks of PCs for baseline vs. SST IN activation at P10 (left). Recordings were subsampled to 360 neurons (200 repeats). Thin lines represent individual mice, thick lines represent the mean. No significant difference between baseline and SST IN activation (Early: β = -0.097, p = 0.125; Intermediate: β = 0.098, p = 0.181). (H) Same as G, but fraction of shared variance explained by descending ranks of shared variance components (left). No significant difference between baseline and SST IN activation (Early: β = -0.083, p = 0.130; Intermediate: β = 0.135, p = 0.043). Ratio of SST IN activation and baseline fractional shared variance spectra (right). (I) Decay exponent α estimated over the stable intermediate range of k = 20-60 for both PCA and SVCA spectra. No significant main effect of Condition on α (Wald χ^2^ = 2.00, p = 0.157) or Method (Wald χ^2^ = 2.96, p = 0.085).

We next tested whether SST IN activation would alter the structure of population activity. SST IN activation did not significantly alter the fractional variance eigenspectrum at either early or intermediate dimensions (**Figure 3G**), and only minimally altered the fractional shared variance spectrum (**Figure 3H**) at P10. Consistent with this, the estimated decay exponent (α) over an intermediate range of dimensions for both PCA and SVCA eigenspectra did not significantly change between baseline and SST activation at P10 (**Figure 3I**). Interestingly, SST IN activation did produce a significant shift in fractional variance at P14, with a significant reduction in early dimensions in the PCA eigenspectrum and significant effects at both early and intermediate dimensions in the SVCA spectrum (**Figure S5E**). The emergence of this effect at P14 but not P10 suggests a dependence on circuit maturation state. Thus, at P10, before the network desynchronizes, SST INs can suppress SNEs without producing a detectable shift in population structure, consistent with the idea that reducing network activity alone is insufficient to shift the distribution of variance across dimensions, and that this ability is specific to PV INs.

### Differential gene expression between P8 and P11 drives unique functional specialization of PV and SST interneurons

We next investigated the molecular pathways underlying MGE IN maturation during the few days of postnatal development when network activity undergoes its dramatic transformation to better understand why PV INs might play a more protagonist role than SST INs and to explore whether cell type-specific changes in gene expression across relevant pathways might explain the circuit level functional differences. To address this, we performed single nucleus RNA-sequencing (snRNA-seq) in S1 at P9 and P11 (**Figures S6A-S6E**). We also reanalyzed a rich snRNA-seq dataset of primary visual cortex (V1) from Gao et al.^28^ to compare the transcriptional programs of PV and SST INs at this critical age (**Figure 4A**).

**Figure 4.**
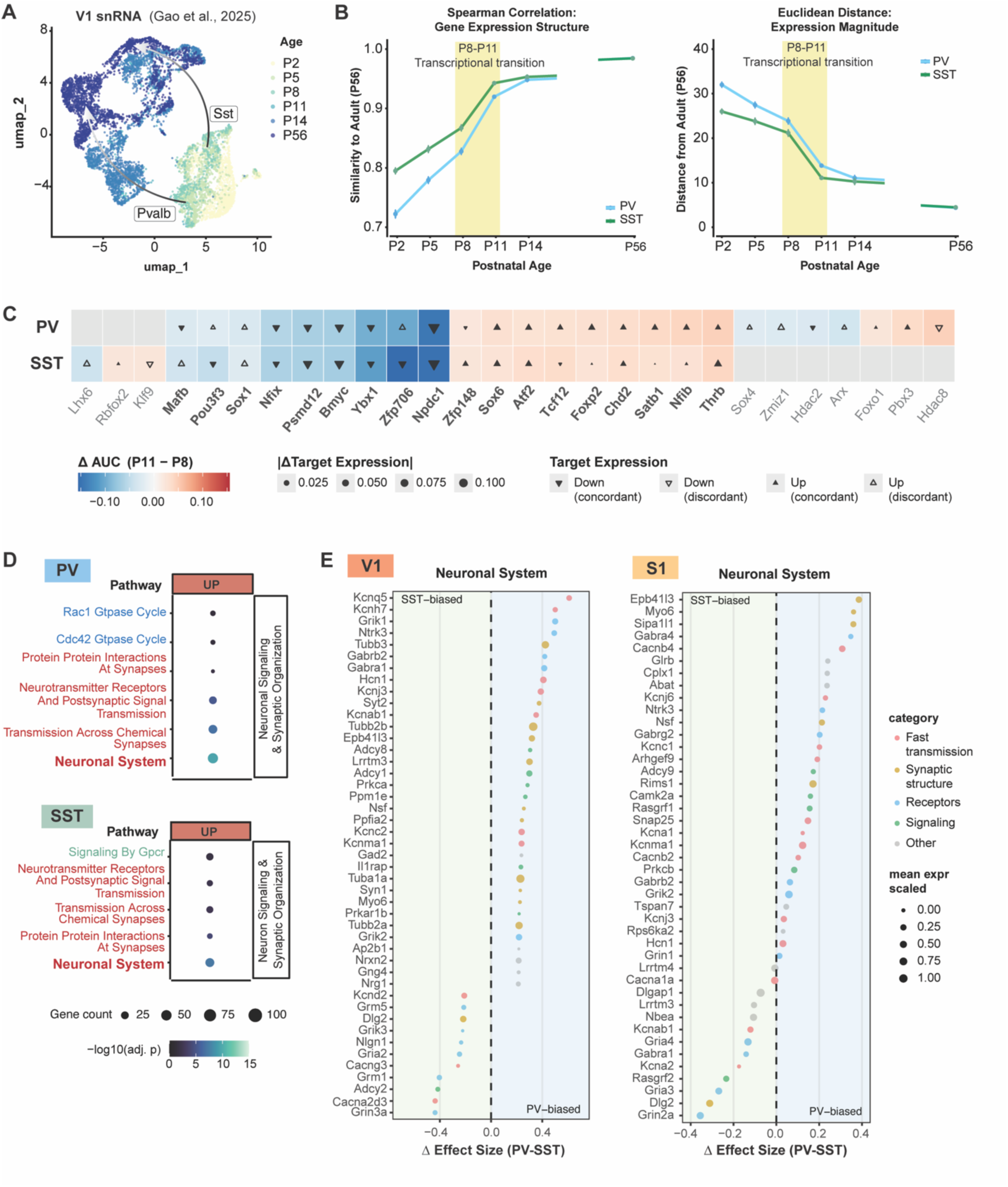
Differential gene expression within shared MGE pathways drives functional specialization of PV interneurons. (A) UMAP of PV and SST INs at different postnatal ages. Data from Gao et al., 2025^28^. (B) Left: Similarity of gene expression structure to mature (P56) state (Spearman correlation) of PV (blue) and SST (green) INs. There is an abrupt increase in similarity between P8 and P11 for both cell types (0.094 in PV, 0.075 in SST across 50 bootstrap iterations). Right: Distance of gene expression magnitude from mature state (Euclidean distance) of PV and SST INs. There is a sharp decrease in distance between P8 and P11 for both cell types (-9.81 for PV, -10.10 for SST across 50 iterations). (C) Heatmap showing the change in transcription factor (TF) regulon activity for the differentially active (Wilcoxon rank-sum test, p < 0.05) regulons in PV and SST INs between P8 and P11. TFs are grouped by specificity: PV-only, shared, and SST-only. Triangle direction indicates whether the mean expression of each TF’s target genes increased or decreased between P8 and P11; filled triangles denote concordance between TF activity and target gene expression changes. (D) Top pathways from over-representation analysis performed using MsigDB Reactome gene sets for PV INs (top) and SST INs (bottom). Upregulated pathways are associated with changes in expression magnitude and pathways associated with neuronal signaling and synapse organization. (E) Relative PV-enriched and SST-enriched genes in the *Neuronal System* Reactome pathway for V1 (left) and S1 (right). Top effect size plotted for each cell type.

We first compared the maturation trajectories of PV and SST INs using pseudobulk gene expression profiles across developmental timepoints relative to mature (P56) reference profiles (see Methods). Maturation was quantified as convergence towards the adult transcriptional state using two complementary metrics, Spearman correlation and Euclidean distance. These metrics measure, respectively, how close the relative ordering of genes is to the mature state at P56 (gene expression hierarchy), and how far expression levels are from the same mature state (gene expression magnitude), which we can use to infer maturational state. Across both metrics, both PV and SST INs exhibited a gradual developmental maturation toward the P56 state (**Figure 4B**). The most pronounced change, however, occurred between P8 and P11, identifying this window as a major transition for transcriptional maturation. Spearman correlation increased sharply over this interval, with the P8-P11 transition accounting for 36.0% of the total increase in transcriptional similarity between P2 and P56 in PV INs and 39.6% of the total increase in SST INs, indicating rapid alignment of gene expression hierarchy with the mature state. Euclidean distance decreased in parallel during the P8-P11 interval, accounting for 36.0% of the total reduction in transcriptional distance from P2 to P56 in PV INs and 46.5% of the total reduction in SST INs, reflecting substantial convergence in gene expression magnitude (**Figure 4B**). Thus, both classes of MGE INs experience a remarkable degree of transcriptional maturation during a narrow developmental window that coincides with the postnatal period in which we observed dramatic changes in network synchrony and population geometry.

We next investigated whether PV and SST INs differ in their transcriptional regulatory mechanisms. We performed gene regulatory network (GRN) analysis on each cell type and assessed changes in transcription factor (TF) regulon activity between P8 and P11, focusing on TFs showing significant up- or downregulation. The majority of significant regulons were shared between PV and SST INs (**Figure 4C**). The shared transcriptional regulatory programs identified by GRN analysis might suggest that PV and SST INs experience similar functional changes during this period. To probe the functional significance of the transcriptomic changes at P8-P11, we performed pathway enrichment analysis. Genes were classified based on their contribution to changes in gene expression hierarchy (gene program) or expression magnitude (magnitude). Consistent with the convergent regulatory programs, PV and SST INs showed similar pathway-level changes (**Figure 4D**). In both cell types, maturation was characterized by upregulation of pathways associated with neuronal signaling and synapse organization, alongside downregulation of pathways associated with protein translation, RNA processing, and genome maintenance (**Figure S7A**). Fewer pathways were driven by changes in gene expression hierarchy alone (**Figure S7A**), suggesting that while large-scale reorganization of gene expression contributes to maturation, it is not the dominant driver at the pathway level; rather, maturation reflects quantitative scaling of gene expression within established programs.

Given this pathway-level similarity, we next asked whether differences between PV and SST INs might emerge at the level of individual genes within shared pathways. Because the *Neuronal System* pathway exhibited both the strongest enrichment and the largest gene set for both PV and SST INs among the pathways commonly upregulated during the transcriptional maturation window, we focused on gene-level contributions within this pathway. PV INs preferentially enriched for genes associated with fast synaptic transmission, including ion channels, synaptic vesicle components, and ligand-gated receptors. In contrast, SST-biased genes were predominantly associated with postsynaptic signaling and dendritic integration, including glutamate receptors, metabotropic receptors, and synaptic scaffolding proteins (**Figure 4E**), a profile consistent with the preferential upregulation of the *GPCR Signaling* pathway in SST INs (**Figure 4D**).

To assess whether these cell type-specific gene expression patterns generalize across sensory cortical regions, we examined PV and SST INs in our own snRNA-seq dataset obtained from S1 and spanning a similar developmental window (P9-P11) (**Figures S6A-S6D**). We discovered that PV INs in S1 showed the same preferential expression of genes associated with synaptic transmission relative to SST INs (**Figure 4E**), indicating that the differential implementation of molecular programs across MGE subtypes is conserved across primary sensory cortical areas.

Thus, we describe a two-tiered organization of MGE IN maturation during the P8-P11 transition. At the regulatory and pathway level, PV and SST INs execute a largely shared transcriptional program, engaging overlapping GRNs and converging on the same broad functional categories. However, within these shared pathways, the two cell types diverge in their specific gene-level implementations, with PV INs elaborating the machinery for fast, precise synaptic output and SST INs focusing on modulatory signaling. This pattern, which is conserved across primary sensory cortical areas, describes a common maturation framework that is differentially tuned, allowing for progressive specialization of each IN subtype.

### Sensory representation is correlated with dimensionality

The rapid developmental transformation of the cortical network to a higher-dimensional regime is thought to result in a sparse and distributed coding scheme^16,17,20^. Given that PV INs are critical for this transition, we next sought to disrupt PV INs to test whether perturbations in the network impact how the S1 population represents sensory information. Instead of brief optogenetic silencing, we triggered programmed cell death in a small fraction of PV INs to permanently remove them from the network after P9. This manipulation is clinically relevant, as a reduced density of PV INs has been reported in human brain tissue from cases diagnosed with Fragile X syndrome^50^ and in several mouse models of autism spectrum disorder (ASD)^51^, and is likely caused by excess developmental cell death^47^.

We selectively expressed the cell death protein Caspase-3 in PV INs by injecting the Cre-dependent rAAV5-Flex-taCasp3 into S1 at P9 (**Figure 5A**). This resulted in a ∼20% decrease in the density of PV INs in S1 of the injected hemisphere relative to contralateral S1 (**Figures S8A** and **S8B**), which matches the reduced PV IN density reported in *Fmr1* and *Shank3* mutant mice^47,50,52–54^. This modest reduction in PV IN density resulted in a trend toward elevated pairwise correlations in the Caspase-3 group at P14 (**Figure 5B**), although the mean PCA eigenspectrum did not show a clear change (**Figure 5C**). This small effect on spontaneous activity suggests that the cortical network is somewhat resilient to mild PV IN perturbations.

**Figure 5.**
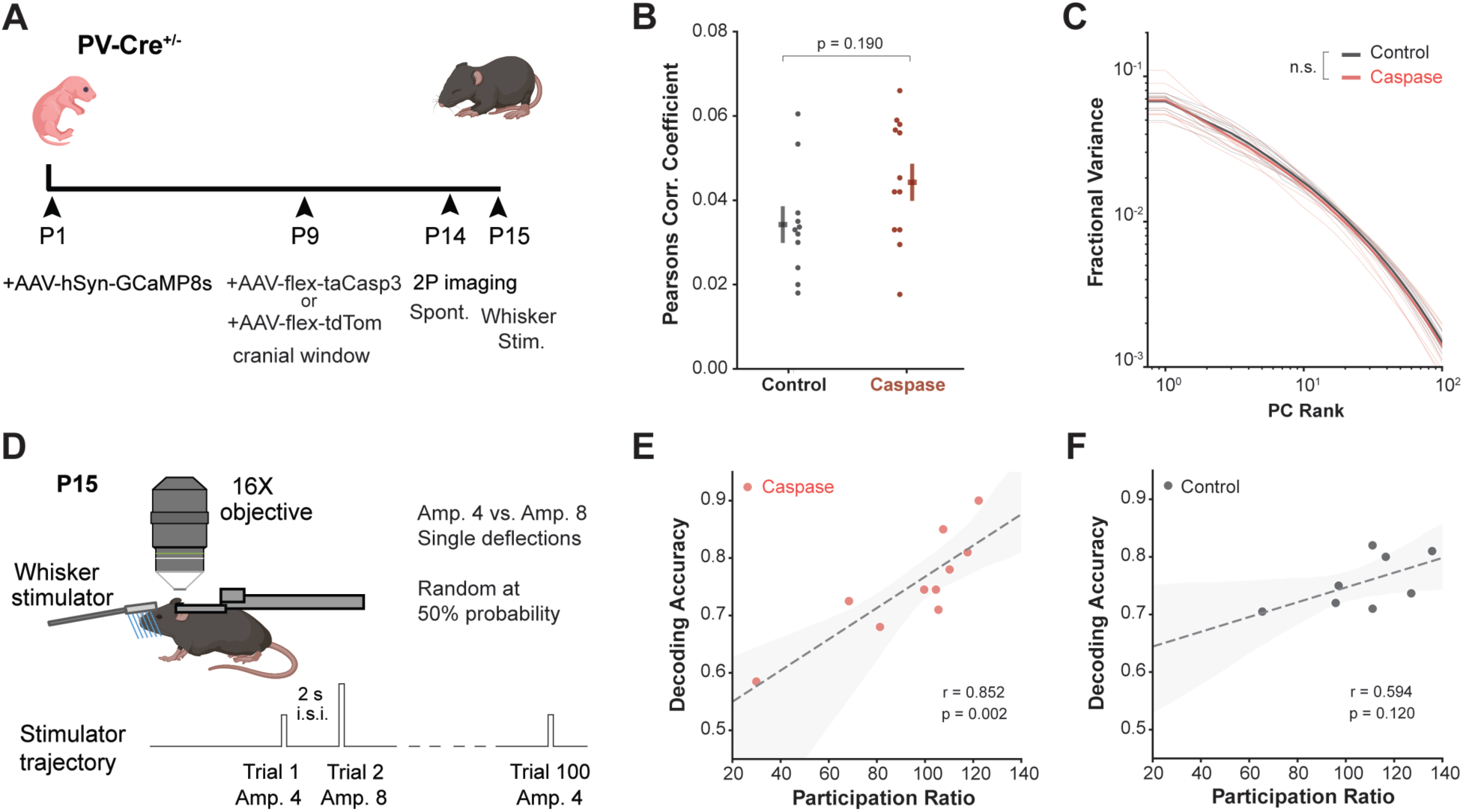
Population dimensionality is correlated with stimulus decoding in Caspase-3 mice. (A) Experimental design for viral-mediated ablation of PV INs. (B) Pearson correlation coefficient (PCC) of neuron pairs. No significant main effect of Experimental Condition (p = 0.190). (C) Fraction of variance explained by descending ranks of PCs plotted on a log-log scale for control mice vs. Caspase-3 mice at P14. Recordings were subsampled to 360 neurons (200 repeats). Thin lines represent individual mice, thick lines represent the mean. No significant difference between control and Caspase-3 groups at early or intermediate ranks (Early: β = - 0.038, p = 0.441; Intermediate: β = -0.088, p = 0.149). (D) Experimental design for whisker stimulation assay at P15. (E) Stimulus-period participation ratio is significantly correlated with decoding accuracy (high d’ neurons, top 20%) for Caspase-3 group. Points represent individual mice. Linear fit (dashed gray line) with 95% CI shown (shaded gray). Pearson r = 0.852, p = 0.002. (F) Stimulus-period participation ratio is not significantly correlated with decoding accuracy (high d’ neurons, top 20%) for control group. Points represent individual mice. Linear fit (dashed gray line) with 95% CI shown (shaded gray). Pearson r = 0.594, p = 0.120.

To assess whether PV cell loss affects sensory encoding, we performed a whisker stimulation assay consisting of 100 trials of brief (50 ms) single-whisker deflections with two different stimulus amplitudes presented in random order (see Methods) (**Figure 5D**). We trained a linear decoder to classify stimulus identity from population activity in S1 using the most stimulus-selective neurons (top 20% by d’). Decoding accuracy was similar between control and Caspase-3 groups (**Figure S8C**), indicating that stimulus information is preserved with a modest reduction in PV IN density. Importantly, we observed a much stronger correlation between population dimensionality during the stimulation period and decoding accuracy in the Caspase-3 group (**Figure 5E**), though not in the control group (**Figure 5F**). Although population dimensionality spanned a similar range across groups, decoding accuracy in the Caspase-3 group covaried more strongly with dimensionality, suggesting that PV IN ablation alters stimulus encoding in proportion to the degree of dimensionality reduction.

### A multi-cell type neural circuit model in which PV neurons are progressively integrated recapitulates developmental changes in neural activity

To better understand the link between developmental changes in cortical circuitry and neural activity, we introduce a biologically realistic, multi-cell-type neural circuit model that captures changes in neural activity regimes between P9 and P14. The recurrent circuit model consists of populations of Exc, PV, and SST neurons, with recurrent connectivity based on the experimental literature^55–57^ (**Figure 6A**), and with intrinsic and extrinsic parameters that evolve with developmental changes also as previously described^58–61^ (**Figures 6B, 6C,** and **S9A-J**).

**Figure 6.**
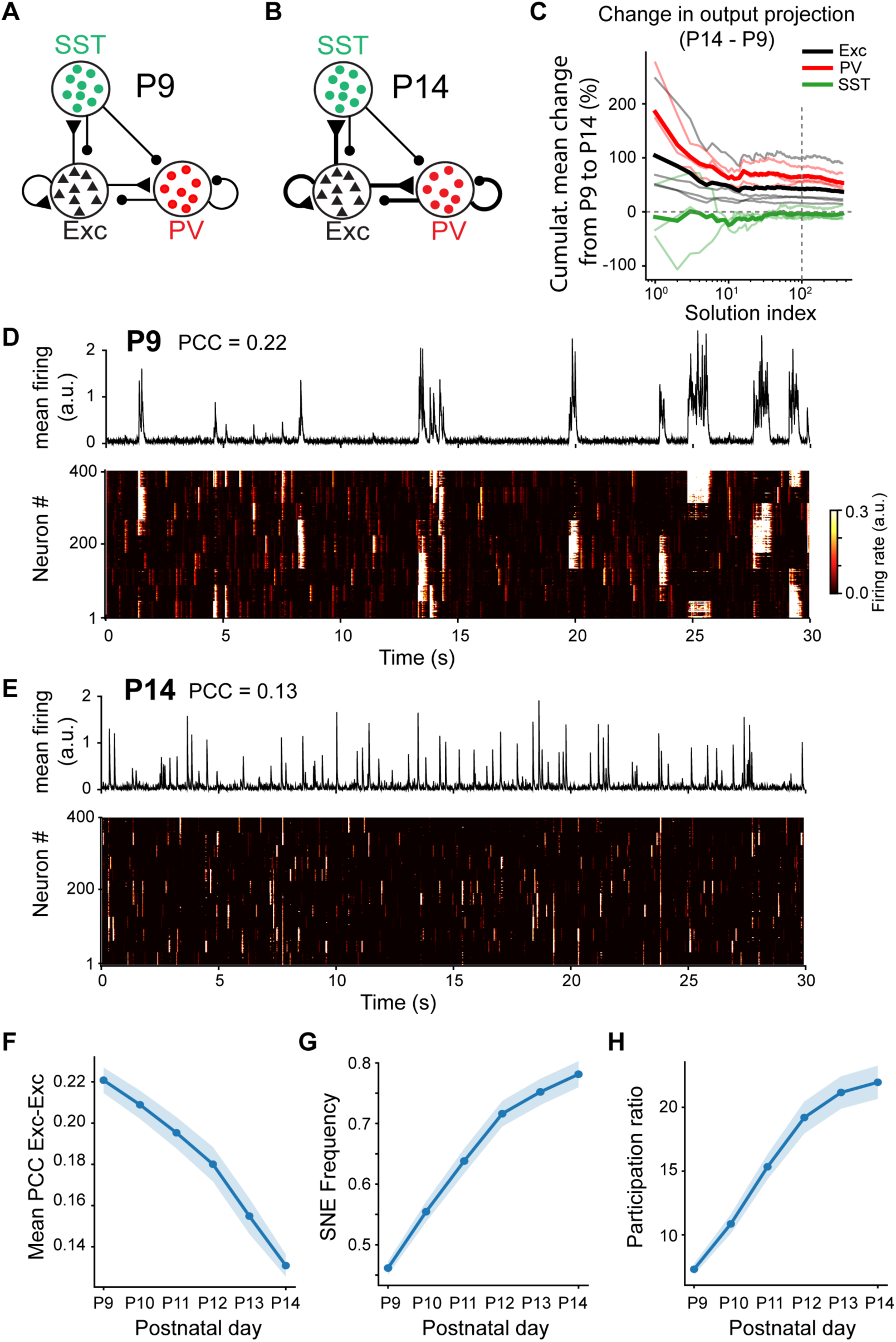
The developmental changes in neural activity are recapitulated by a mechanistic model that increases Excitatory and PV parameters. (A) The cortex at P9 was modeled with a recurrent multi-cell-type population circuit of 400 Exc, 40 PV, 40 SST neurons. (B) Same as A, but with thicker lines representing the parameters that are required to increase in development: synaptic connections and intrinsic threshold (the latter corresponding to a decrease in excitability represented by a thicker circle surrounding a population). (C) Top 500 solutions sorted based on biological alignment with development and neural activity statistics for the model parameters’ change from P9 to P14 for Exc (black), PV (red), SST (green) neurons. The solutions are cumulatively averaged. Individual thin lines correspond to single parameters, while each thick line corresponds to the average of parameters belonging to the same population. The dashed vertical line represents the top 100 solutions that we averaged over for panels F-H. (D) Top: Average firing rate (arbitrary units) of the model at P9. Bottom: The corresponding activity of single Exc neurons as a function of time. (E) Same as D, but for the P14 circuit. (F-H) Change in development for three main neural activity summary statistics: PCC between Exc neurons (F), SNE frequency (G), and Participation ratio (H). The solid and shaded lines represent, respectively, the mean and the standard error over the first 100 parameter sets, sorted as in C.

We sought to reproduce the slow, second-scale dynamics of SNEs while generating these events intrinsically within a recurrent, single-neuron-resolution, multi-cell-type circuit model. Existing models of developmental desynchronization^30,32–35,37,38^ incorporate only a subset of these network properties while recurrent, single-neuron-resolution, multi-cell-type circuit models that focus on the adult mouse brain and support intrinsically generated fluctuations^39,40^ do not reproduce the second timescales observed in the data.

Our model successfully reproduced the key features of network activity observed in calcium imaging data: at P9, neural dynamics exhibit pronounced population-wide fluctuations, characterized by alternating low- and high-activity periods with intermediate PCCs (**Figure 6D**); while at P14 the dynamics exhibit a more desynchronized activity pattern with heterogeneous firing across neurons and lower PCCs (**Figure 6E**). Synaptic connections were structured in a way that supported clustered patterns of activity^43,62^, in which only a subset of neurons would have elevated firing rate (local SNEs). Absent other features, such as periods of silence, this dynamic structure would exhibit unrealistically low firing rates during development^43^, while the simple alternation between periods of silence and global SNEs would exhibit unrealistically high correlations (**Figure S9K**). Our network solves this conundrum by alternating periods of silence with periods of local SNEs, which induces low but non-negligible correlations (PCC ∼0.1) compatible with the experimental data. Clustering in the excitatory connections results in clustered activity bursts and, consequently, in low PCC (**Figure S9K**), even in the presence of strong fluctuations in the average population rate (as measured by a bimodality index; see Methods).

We sought parameter sets such that the solutions would reproduce key neural statistics^63,64^ and developmental trajectories. Specifically, we required the model to reproduce developmental changes in average PCCs, SNE frequency, participation ratio, and strong fluctuations in the average rate. Additionally, we required the parameters to align as closely as possible with known developmental changes (**Figure 6C**). Although the existence of solutions exhibiting the observed changes in structure and neural dynamics was not guaranteed *a priori*, our network simulations revealed that such solutions exist across a broad range of parameter values.

Averaging across the top 100 biologically aligned solutions confirmed that the circuit model exhibited during development: a progressive decrease in pairwise correlations between excitatory neurons (**Figure 6F**); an increase in the frequency of SNEs (**Figure 6G**); and an increase in participation ratio (**Figure 6H**). Finally, the model’s average excitatory firing rate exhibited a bimodal distribution at early developmental stages, corresponding to alternating low- and high-activity states, and this bimodality progressively weakened from P9 to P14 (**Figure S9L**). These results show that a recurrent Exc–PV–SST circuit in which excitatory and PV parameters progressively strengthen during development can reproduce the experimentally observed changes.

### The model predicts changes in PCC elicited by activation of PV INs

We then asked whether our circuit model could predict changes in PCCs caused by cell-type-specific optogenetic stimulation. We considered the top biologically aligned model solutions (**Figure 6**), which were not specifically selected to reproduce responses to optogenetic perturbations. At P9, the prediction matches our data, as PV, but not SST, stimulation reduced spontaneous pairwise correlations (**Figure 7A** and **7B**), as observed experimentally (**Figures 2** and **3**). By P14, both model and data show only weak effects of either manipulation (**Figures 7A** and **2**).

**Figure 7.**
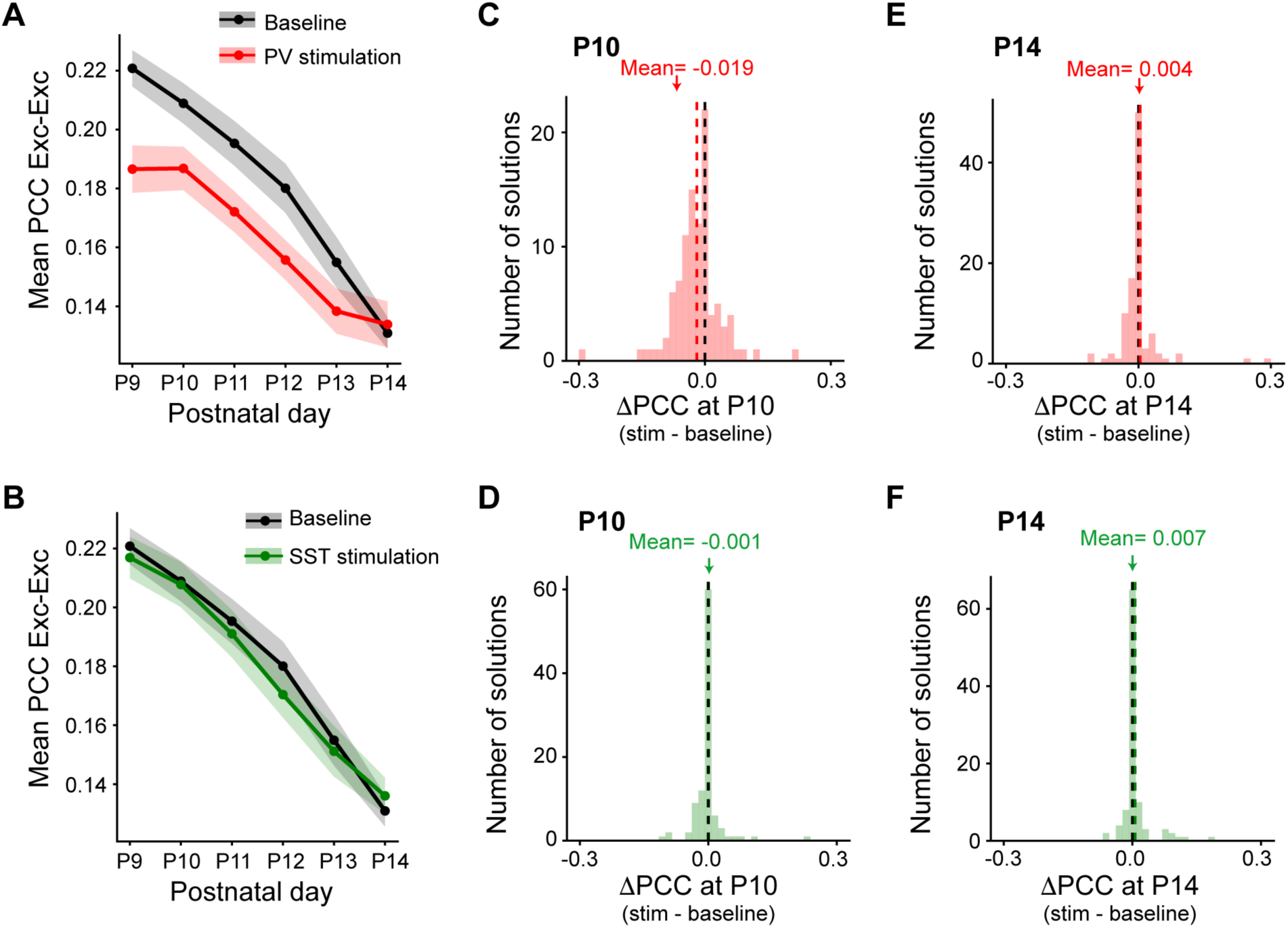
The mechanistic model predicts optogenetic changes in PCC that align with the experiments. (A) Average across the top 100 solutions of the sorting introduced in Figure 6C of PCC between excitatory neurons across development for the baseline results and the predicted ones for the optogenetic stimulation of PV neurons. (B) Same as A, but the optogenetic stimulation was for SST neurons. (C) Histogram of the predicted change in PCC at P10 before and after optogenetic stimulation of PV neurons. (D) Same as C, but for the SST optogenetic stimulation. (E and F) Same as C and D, respectively, but at P14.

The optogenetic prediction was robust across solutions. At P10, the distribution of PCC induced by PV stimulation was shifted toward negative values (**Figure 7C**), whereas the distribution for SST stimulation remained centered near zero (**Figure 7D**). By P14, both distributions were concentrated around zero, indicating that the effect of optogenetic perturbations on network correlations diminished with development (**Figures 7E** and **7F**). These results were unchanged as the number of retained parameter sets increased (**Figures S10A** and **S10B**). Across the top 100 parameter sets, the optogenetically induced changes in PCC covaried with changes in network properties in the participation ratio and in SNE occupancy, defined as the fraction of time spent in synchronous network events (**Figures S10C**). Together, these results suggest that PV IN modulation influences network correlations by reshaping the temporal organization of collective activity.

## DISCUSSION

The remarkable transformation of cortical network activity that takes place during the second postnatal week in rodents (corresponding to the time of birth in humans), has been extensively characterized phenomenologically^2,4,65,66^. Over a period of just a few days, from P9 to P13, population activity becomes more continuous, with neurons firing sparsely, a regime that has been predicted to enable a greater coding capacity. The goals of this study were three-fold: first, to investigate the circuit and molecular mechanisms that bring about this critical developmental milestone; second, to better understand the relationship between population dynamics and information encoding; and finally, to develop a biologically plausible computational model of developmental network activity that could assist in future studies (e.g., by making predictions about potential mechanisms, etc.).

Our longitudinal recordings of spontaneous activity between P9 and P14, combined with IN perturbations and single-cell transcriptomics, reveal five key findings: 1) The developmental expansion of dimensionality reflects a redistribution of fractional variance from a few leading dimensions toward a broader, more uniform distribution across the population. 2) Acute optogenetic activation of PV INs but not SST INs at P10 shifts fractional variance towards a more uniform distribution, implicating PV IN activity as a driver of the redistribution of fractional variance that accompanies dimensionality expansion during development. 3) Between P8 and P11, PV and SST INs undergo rapid transcriptional maturation but diverge in the regulation of genes associated with fast synaptic transmission, which are selectively enriched in PV INs, providing a molecular basis for their distinct functional role. 4) Networks with higher dimensionality support greater stimulus encoding capacity, linking developmental circuit-level reorganization to the cortex’s ability to represent sensory information. 5) A new computational model reproduces developmental trajectories of population activity and predicts the privileged role of PV INs in driving network desynchronization.

The developmental reorganization we describe suggests that circuit maturation does not merely reflect the elaboration of existing structures or the loss of coordination but instead arises from a fundamental redistribution of fractional variance from a few dominant, synchronous modes into many weaker dimensions. The shift towards a more distributed eigenspectrum and fractional shared variance spectrum across postnatal development is a progression toward the higher-dimensional, structured spontaneous activity reported in adult primary sensory cortex^21^.

Our optogenetics results extend previous observations using activation of a broad IN population driven by pan-GABAergic promoters such as *Gad2* or *Gad67*^30,48^ by demonstrating the protagonist role of PV INs over SST INs. The acute activation of PV INs significantly decorrelated activity at P10 in a distance-dependent manner, consistent with previous work^30,31^. Although acute optogenetic activation of PV INs drove decorrelation, it did not completely recapitulate the redistribution of fractional variance observed in development. This could be attributed to limitations in our experimental approach, such as the fact that ChRmine was only expressed in a subset of PV INs (likely ∼30%^47^) or that our laser scanning was restricted to only a small volume of S1. Another possibility is that because PV INs are not fully mature, their synaptic connections, both to other PV INs and to the Exc population, have not yet reached full strength, further limiting the effect of acute PV IN activation at the early timepoint of P10. Indeed, although PV INs are functionally connected to Exc neurons at P10^47^, the peak of PV synaptogenesis occurs around P10-P12^29^.

It is likely that other maturational factors are necessary as PV activation did not fully recapitulate the developmental reorganization of network activity. Besides PV INs, the significant reduction in the intrinsic excitability of the Exc population in the network may play a role. Indeed, the resting membrane potential of cortical pyramidal neurons decreases between P6-8 and P13-16^67^, presumably because of changes in the expression of leak conductances, including I_h_, as previously proposed^5^. The comparatively depolarized state of the P10 network could limit how far acute PV IN activation alone can push dimensionality. Other developmental changes, such as the transition from transient excitatory corticothalamic amplification to corticothalamic recruitment of inhibition^15^, could also contribute.

Interestingly, PV IN activation also significantly increased fractional shared variance in the intermediate dimensions at P14, though with a smaller effect size than at P10 (**Figure S3F**), despite no significant change in the PCA eigenspectrum. This is consistent with acute PV IN activation having a reduced capacity to further reorganize an already-redistributed population structure, while retaining a residual capacity to modulate the reliability of activity within the existing framework. Likewise, the increased ability of SST INs to shift fractional shared variance at P14 (**Figure S5F**) could reflect a dependence on circuit maturation; SST INs’ modulatory action may require a mature dimensional structure that is not present until later in development, potentially as a consequence of PV-mediated redistribution.

Our findings add to a proposed framework in which GABAergic INs sequentially control network activity during development^4,48,68^. SST-expressing hub cells orchestrate network synchrony early in postnatal development^69,70^ while PV INs become the dominant influence during the second postnatal week, undergoing a period of transcriptional maturation that gives them the ability to drive an expansion of dimensionality. This is consistent with prior studies demonstrating a role for SST INs in PV circuit maturation during the first postnatal week in both deep and superficial layers^31,61^.

A protagonist role for PV INs in desynchronization is consistent with the timing of their maturation. They assume their chemical and electrophysiological identity at P9-P10^24,60,71^, as they begin to express *Pvalb* and *Kv3.1b*, the potassium channel that allows them to be fast-spiking^58,72,73^. There is also a strengthening of reciprocal glutamatergic drive onto PV INs themselves throughout this time window^74^. The differential effects of PV and SST INs on correlated activity and dimensional structure may reflect cell type-specific implementation of shared maturational programs acting on already established circuit architecture. PV INs, which target the perisomatic region of pyramidal cells^75,76^, preferentially express genes supporting fast synaptic transmission during the P8-P11 transition window, while SST INs, which preferentially make synapses onto apical dendrites^77,78^, exhibit a bias towards modulatory signaling components.

The covariation between stimulus encoding and network dimensionality in the Caspase-3 mice, whose degree of PV IN loss was variable, may have clinical relevance for neurodevelopmental disorders associated with PV IN deficits. For example, the overall ∼20% reduction in PV IN numbers we achieved with Caspase-3 expression is similar to that reported in the *Fmr1* knockout mouse model of Fragile X syndrome^47,52,53^, which demonstrates a delayed desynchronization of network activity^79,80^. As in models of Fragile X syndrome^47,53^, the degree of reduction in PV IN density in the Caspase-3 group was variable across animals. This variability and its corresponding relationship with decoding accuracy suggests that a spectrum of PV IN density loss may produce correspondingly graded reductions in network dimensionality and stimulus encoding capacity, providing a potential link between the degree of PV IN deficit and the variability of behavioral symptoms such as sensory hypersensitivity and cognitive impairment. Future work examining how dimensionality evolves during postnatal development across neurodevelopmental disorders, both during spontaneous activity and during sensory stimulation, could directly test this hypothesis.

It is difficult for the data alone to isolate which of the maturational changes drives the reorganization of activity: Exc, PV, and SST populations are embedded in recurrent loops, so the effect of strengthening a synapse or shifting an excitability threshold depends on every other concurrent change. To disentangle these interacting factors and establish a mechanistic link between the developmental reorganization of cortical activity and maturation of the underlying circuitry, we introduced a biologically realistic multi-cell-type recurrent circuit model. Our model was designed to overcome limitations of previous models of developmental desynchronization^30,32–35,37,38^, as described in the Introduction. It is, to our knowledge, the first to combine recurrent connectivity, single-neuron resolution, and three neuronal cell types, while generating seconds-scale synchronous events intrinsically that match pairwise correlations measured in vivo. A broad range of model solutions satisfying biologically motivated developmental constraints reproduced the principal features of this transition, including reduced Exc-Exc pairwise correlations, increased SNE frequency, and increased activity dimensionality. These results suggest that the developmental reorganization of recurrent Exc and PV circuitry can generate a transition from a network dominated by large, population-wide fluctuations to one characterized by more frequent but spatially restricted events. Importantly, the model does not require external inputs to drive synchronous events, suggesting that the slow, seconds-scale fluctuations observed experimentally can emerge intrinsically from recurrent interactions among distinct neuronal populations. The model also provides a mechanistic explanation for why relatively weak average pairwise correlations, typically associated with stationary global population activity^81,82^, here coexist with strong fluctuations^83,84^: clustered excitatory connectivity promotes local synchronous events while preventing global synchronization. Finally, we recapitulated the developmental specificity of PV-mediated modulation observed experimentally, predicting that PV activation reduces excitatory correlations early in development while having little effect at later stages. This modulation was associated with changes in SNE occupancy, suggesting that PV neurons regulate correlations by reshaping the temporal organization of the collective network. Our model provides a mechanistic link between circuit maturation, the emergence of higher-dimensional activity, and the developmental loss of global synchrony, and it opens new avenues for hypothesis generation that will undoubtedly motivate future in vivo experiments.

## ACKNOWLEDGEMENTS

We are grateful to Claudia Nguyen, Elisa Fazzari, and Daria Azizad for advice on single-nucleus RNA sequencing experiments and data processing, to Aparna Bhaduri for guidance on transcriptomics analysis, and to Dean Buonomano, John Zhou and Katheryn Zhou for early discussions on the computational model. We thank Peyman Golshani for feedback on the manuscript. This work was supported by the following grants: R01HD108370 (NIH-NICHD), R01HD054453 (NIH-NICHD), and a pilot grant (# 01POR2024) from UCLA’s Center for Autism Research and Treatment (CART) and the Clinical and Translational Science Institute (CTSI) awarded to C.P.-C.; R21EY035064 (NIH-NEI) awarded to M.D.; the Training Grant in Neurobehavioral Genetics (T32NS048004; NIH-NINDS), F30HD117619 (NIH-NICHD), and the UCLA-Caltech Medical Scientist Training Program (T32 GM008042 and T32 GM152342; NIH-NIGMS) to M.W.W.

## DECLARATION OF INTERESTS

The authors declare no competing interests.

## DECLARATION OF GENERATIVE AI AND AI-ASSISTED TECHNOLOGIES IN THE MANUSCRIPT PREPARATION PROCESS

During the preparation of this work, the authors used Claude to edit the text for clarity and grammar. After using this tool, the authors reviewed and edited the content as needed and take full responsibility for the content of the published article.

## EXPERIMENTAL MODEL

### Mice Lines

All experiments followed the U.S. National Institutes of Health guidelines for animal research, under an animal use protocol (ARC #2007-035) approved by the Chancellor’s Animal Research Committee and Office for Animal Research Oversight at the University of California, Los Angeles. All mice were housed in a vivarium with a 12/12 h light/dark cycle and experiments were performed during the light cycle. Mice of both sexes were used in this study, and all were on the C57BL/6 background strain.

Mouse lines were obtained from Jackson laboratory: *Nkx2.1*-Cre mice (JAX 008661), *SST*-Flp (JAX 031629), PV-Cre (JAX 008069), and Ai14 (JAX 007914).

## METHOD DETAILS

### Viral injections

Stereotaxic viral injections were done using glass micropipettes (Sutter Instrument, 1.5 mm outer diameter, 0.86 mm inner diameter) through burr holes at P0-P1. Briefly, prior to viral injection, custom micropipettes were made by pulling the micropipette using a dual-stage glass pipette puller (PC-100, Narishige), such that the length of the tapered tip was 6-8 mm and the outer diameter was 10-15 µm. All mice were anesthetized with isoflurane (5% induction, 1-1.5% maintenance via a nose cone, v/v) and placed in a stereotaxic frame with rubber ear bars. Pups were allowed to recover on a warm water circulation blanket (T/Pump, Gaymar Stryker) before being returned to the dam. For injections at P0-P1, we made a small skin incision on the scalp over the right hemisphere under sterile conditions and drilled a small burr hole over the right S1 cortex using a dental drill, as previously described (He, Arroyo et al 2018). Multiple injections were performed using a Picospritzer (General Valve, 30 pulses of 6 ms, 30 psi) for a final volume of 250-350 nL. For injections at P9 (Caspase-3) we injected the rAAV at the time of the cranial window surgery at multiple locations.

We used the following viruses: pGP-AAV-syn-jGCaMP8s-WPRE (Addgene 162374-AAV1), AAV1-CAG-FLEX-jGCaMP8s-WPRE-SV40 (Addgene 162380-AAV1), pAAV-nEF-Con/Foff 2.0-ChRmine-oScarlet (Addgene 137161-AAV8), pAAV-nEF-Coff/Fon-ChRmine-oScarlet (Addgene 137160-AAV8), pAAV-Ef1a-Coff/Fon-mCherry (Addgene 137134-AAV8), pAAV-Ef1a-fDIO-GCaMP6s (Addgene 105714-AAV8), and pAAV-flex-taCasp3-TEVp (Addgene 45580-AAV5).

### Cranial windows

Pups were anesthetized with isoflurane (3.5% induction, 1.5–2% maintenance via a nose cone, v/v) and secured in a stereotaxic frame with rubber ear bars. A 3.5-4 mm diameter craniotomy was performed over the right S1 cortex, under sterile conditions, by removing the skull without disturbing the pia, as described previously^85^. The craniotomy was then covered with a 4 mm glass coverslip and secured by cyanoacrylate glue and dental cement. A custom titanium head bar was attached to the skull caudal to the window with dental cement to secure the animal to the microscope stage. Within 1 h after surgery, the pups appeared fully recovered from the effects of anesthesia and could be returned to their dam. For in vivo imaging at P9, cranial windows were performed on the day before the recording under 1-1.5% isoflurane anesthesia (3.5% induction) and we allowed mice to recover for 4-6 h with their dam prior to calcium imaging.

### In vivo 2-photon calcium imaging in head-restrained mice

Calcium imaging was performed using a DIY Bergamo II scope (Thorlabs) at 15.13 Hz with a 16X objective (Nilon, NA=0.8), which allowed for a large field of view (512 x 512 pixels corresponding to 722 x 722 µm). The microscope was coupled to a Chameleon Ultra II Ti:sapphire laser (Coherent) tuned to 930 nm for calcium imaging (average power at the sample was <90 mW). We used a separate 1,040 nm laser (Fidelity) for optogenetic manipulations.

Calcium imaging was done at a depth of 130-190 μm below the dura for P9 mice and up to 220-250 μm for P14-15 mice, corresponding to L2/3. For all recordings, mice were unanesthetized and placed in a cotton pad and kept at 37°C using the same heating system. Recording sessions lasted <45 min.

### Whisker stimulation protocol

For calcium imaging of sensory-evoked activity, whisker stimulation was achieved by bundling the contralateral whiskers (typically all macrovibrissae of at least 1 cm in length) with soft bone wax and attaching them to a glass capillary coupled to a piezo-actuator (Physik Instrumente). The stimulation protocol consisted of a 30 s baseline followed by whisker stimulations using 50 ms deflections along the anterior–posterior direction, with a 2 s interstimulus interval (ISI).

Deflections of two different amplitudes (A4 and A8) were delivered by changing the voltage applied to the piezo-actuator.

### In vivo 2P optogenetic stimulation

Optogenetic photostimulation of ChRmine-expressing neurons was performed, as previously described (Kourdougli et al., 2023), by scanning a separate 1,040 nm femtosecond-pulsed laser (Fidelity, Coherent) using the same 2P microscope. Laser power was typically ∼100 mW at the objective. We used two different photostimulation protocols. The first consisted of 90s of baseline followed by six 10 s-long pulses with an inter-stimulation interval of 20 s, and the second one consisted of 90s of baseline followed by twenty 1 s-long stimulation pulses, with an I.S.I. of 3 s. The laser beam was controlled by an optical beam shutter controller (SC10, Thorlabs) triggered via custom-built MATLAB code.

### Calcium imaging data analysis

#### Preprocessing

Motion correction and segmentation was performed using Suite2p^86^. Recordings with >400 detected neurons were used for analysis. Raw fluorescence was neuropil-subtracted and normalized using custom Python scripts. Deconvolution was performed with the Suite2p implementation of OASIS. A modified Z-score was used for normalization of both raw and deconvolved traces. Briefly, a quiet baseline period was identified and used to Z-score the neuropil-subtracted fluorescence to normalize while preserving neuronal activity variance:

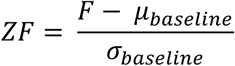

#### Correlations

Pearson correlation coefficients were calculated using deconvolved, non-binarized traces. The entire trace was used. For distance analysis, neurons pairs were binned by their physical distance in 100 μm increments for pairwise correlations.

For correlations versus chance, a subset of 10,000 neuron pairs per recording was used. 50 chance distributions were generated by randomly circularly shifting the trace of 1 randomly chosen neuron and computing the correlation to the other neurons. Traces were binned (to 1s) and z-scored for this calculation.

For mean firing-rate correction, the mean firing-rate proxy (derived from deconvolved activity) was calculated for each neuron pair. Neuron pairs were grouped into bins according to their mean firing rate (bin width = 0.02; maximum = 0.10, based on the firing-rate distribution). To balance observations across recordings, periods, and firing-rate bins, 1000 neuron pairs were randomly sampled without replacement from each recording session, experimental period (Baseline or Stimulus), and firing-rate bin. Pairwise correlations from these sampled neuron pairs were analyzed using a linear mixed-effects model (LMM) with Firing-Rate Bin, Period, and their interaction as fixed effects and Subject (mouse ID) as a random intercept. When the omnibus Wald test identified a significant Firing-Rate Bin × Period interaction, post hoc LMMs were fit separately within each firing-rate bin to compare Condition (Baseline or Stimulus), with Benjamini–Hochberg false discovery rate correction for multiple comparisons. For visualization, pairwise correlations were averaged within each mouse for each firing-rate bin and period, and data are presented as mean ± SEM across mice.

#### Synchronous network events

To detect synchronous network events, we used binarized activity vectors. These were constructed from the deconvolved traces with a threshold of 2 standard deviations above the mean. Binarized activity vectors were randomly circularly shifted to create 1,000 surrogate distributions, and a sum of active cells vector was created for each frame. The 97.5^th^ percentile was used as a statistical threshold. Peaks that were at least 3 frames long were considered SNEs. The onset and offset of a SNE are defined as the first frame above threshold and the first frame below threshold respectively.

#### Coefficient of Variation

Coefficient of variation was defined as the ratio of the standard deviation of the fraction active to the mean of the fraction active, where the fraction active is the percent of cells active in each SNE.

#### Participation ratio

The participation ratio was computed from deconvolved, non-binarized neural activity traces. It was defined as the ratio between the second and the first moments of the spectral density, obtained via eigen decomposition of the covariance matrix of neural activity^46,87^. To ensure robustness to sampling, participation ratio reported was the average of 200 independent subsamples of n=360 neurons (sampled without repeats).

#### Principal components analysis

We computed the eigenspectrum of the covariance matrix of the full population activity (modified Z-scored fluorescence):

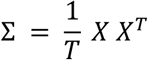

Eigenvalues λ_k_ were obtained via eigendecomposition of the covariance matrix and represent the total variance (signal + noise) in each principal component (PC).

To ensure robustness to sampling, analyses were performed on random subsets of neurons. For each dataset, we drew subsamples of size n=90, 180, and 360 (without replacement), and repeated all analyses across 200 independent subsamples. Results were averaged across the 200 subsamples. n=360 was used for most of the analyses.

Fractional variance is defined as: 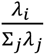

#### Shared variance components analysis

A time-split SVCA, adapted from SVCA^21^, identifies dimensions of neural activity that are reproducible across independent subsets of time.

Data preprocessing: Neural activity traces (modified Z-scored) were mean-centered.

Subsampling: To ensure robustness to sampling, analyses were performed on random subsets of neurons. For each dataset, we drew subsamples of size n=180, 360, and 720 (without replacement), and repeated all analyses across 200 independent subsamples. Results were averaged across the 200 subsamples. N=360 was used for most of the analysis, as n=360 and n=720 give similar results up to ∼100 dimensions.

Temporal splitting for spontaneous activity: Timepoints were divided into non-overlapping contiguous segments of 150 frames (approximately 10s) and segments were randomly assigned into two equal sets. This procedure ensured that estimates of shared variance were based on activity patterns that generalized across time. For each subsample, this splitting procedure was repeated 5 times, and results were averaged across splits.

Temporal splitting for optogenetics: For trial-structured datasets, independent data subsets were defined at the level of trials. For each split, trials were randomly permuted and partitioned into two non-overlapping groups of equal size. Activity from each group was concatenated across trials. This procedure ensured that the two datasets were statistically independent while preserving within-trial temporal structure, providing an estimate of reliability across independent repetitions of the same experimental condition. The random partitioning of trials was repeated 5 times for each subsample, and all quantities were averaged across splits.

Estimation of shared variance components: For each split, the cross-covariance matrix was computed from the two time subsets.

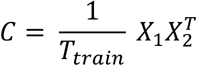

Singular value decomposition (SVD) of C was used to identify pairs of projection vectors that maximize covariance between the two neuron groups. These projections define the shared variance components (SVCs).

Estimation of reliable variance: For each component, reliability was defined as the Pearson correlation between projections of the two time subsets:

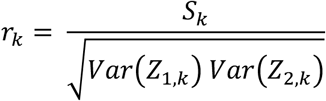

Where *S_k_* is the singular value of the cross-covariance matrix and *Z_1,k_* and *Z_2,k_* are the corresponding projections. This provides an estimate of the fraction of variance that is reproducible across independent temporal samples. Because neurons were not split into independent groups, this measure primarily reflects temporal reliability and may modestly overestimate shared variance compared to fully cross-validated approaches.

#### Signal fraction

To quantify the fraction of variance attributable to reliable activity, we computed:

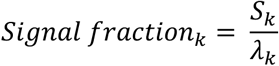

#### Decay exponent

The decay exponent α was obtained by fitting a power law to the eigenspectrum over an intermediate range of components (k = 20-60).

#### PC loadings

To assess how Exc and Inh neurons contribute to each PC relative to their numeric proportion in the population, we computed the inhibitory overrepresentation in PC loadings as a function of PC rank.

#### Decoder

Deconvolved neural population activity was organized into a three-dimensional tensor (neurons x time x trials). To quantify stimulus selectivity of individual neurons, we computed the sensitivity index (d’) using the mean activity across the analysis window. For each neuron, responses to the two stimulus conditions (amplitude 4 and amplitude 8) were separated, and d’ was calculated as:

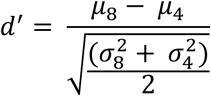

where μ and σ^2^ denote the mean and variance of trial-averaged firing rates for each stimulus condition. Neurons were ranked according to the absolute value of d’. High-selectivity neurons were defined as having |d’| greater than or equal to the 80^th^ percentile of the distribution.

Stimulus identity was decoded using L2-regularized logistic regression. Neural activity from the high-selectivity neurons was reshaped into a feature matrix by concatenating activity across all time bins to produce one feature vector per trial. Features were z-scored using statistics computed from training data only. Decoder performance was evaluated using 4-fold cross-validation to preserve the proportion of stimulus classes in each fold. Classification accuracy was averaged across folds. To estimate chance performance, stimulus labels were randomly permuted 20 times while keeping neural activity unchanged. The complete cross-validation procedure was repeated for each permutation, and the mean of the shuffled accuracy was used as the empirical chance baseline. Decoder performance exceeding the shuffled-label accuracy indicated that population activity contained information about stimulus identity.

To assess the relationship between participation ratio and stimulus encoding capacity, Pearson correlation was computed between participation ratio and decoding accuracy separately for control and caspase-3 groups. Linear fits were computed using ordinary least squares regression for visualization.

### Statistical analysis

Linear mixed-effects models (LMM) were used with appropriate fixed effects (Day, Cell Type, Condition) and with Subject (mouse ID) as a random intercept using the statsmodels package in Python^88^. Residual normality was assessed using the Shapiro-Wilk test and by visual inspection of quantile-quantile (QQ) plots; non-normal data was log-transformed or the rank-inverse normal was used to satisfy normality. The Benjamini-Hochberg procedure was used to correct for false discovery rate. Omnibus Wald test was used to test factors of LMM. Data are presented as mean ± SEM across mice (Figs. 1, 2, 3, and 6).

To test whether interneuron activation differentially affected fractional variance and reliable variance in early versus late dimensions, we averaged each measure across early (k=1–5) and late (k=20–60) dimensions separately for each recording. Dimension ranges were defined based on the crossover point identified in the developmental eigenspectrum comparing P09 and P14, where early dimensions correspond to those losing reliable variance across development and late dimensions correspond to those gaining it.

Variance was log-transformed prior to analysis to better satisfy normality assumptions. For each dimension range, we fit a LMM with condition (Baseline or Stimulus) as a fixed effect and subject (mouse ID) as a random intercept. Residual normality was assessed using the Shapiro-Wilk test. We report the fixed effect coefficient (β) and the p-value for the condition term from each model. To test whether interneuron activation affected reliable variance fraction, we performed an analogous analysis using reliable variance fraction as the outcome variable, averaged over the same dimension ranges.

### RNA-sequencing

#### RNA extraction

Mice were anaesthetized with isoflurane, followed by cardiac perfusion with ice cold choline ACSF containing 132 mM choline, 2.5 mM KCl, 1.25 mM NaH2PO4, 25 mM NaHCO3, 7 mM MgCl2, 0.5 mM CaCl2, and 8 mM D-glucose) as previously described (Suresh et al., 2026). Coronal sections (1 mm thick) containing S1 (A/P -1.0 to - 2.0 mm, M/L 2.0 to 3.0 mm) were collected using a Stainless-Steel Brain Matrices (Zivic instruments) in chilled choline, from which S1 was dissected bilaterally and snap-frozen in liquid nitrogen and stored at -80°C in preparation for single-nuclei RNA-sequencing. Samples were pooled across mice (n=6 each at P9, P11, and P13).

#### Library preparation and sequencing

Nuclei were extracted from frozen S1 samples using the 10X Chromium Nuclei Isolation Kit, counted using a hemacytometer, then resuspended in buffer to achieve a target capture of 10,000 nuclei. Samples were processed using the 10X Genomics Chromium Single Cell 3’ Reagent Kit v4 and assessed for quality control purposes at both the cDNA and library stages using the Agilent 4200 TapeStation System. Single-nuclei libraries were sequenced using the Illumina Novaseq X platform.

#### Data processing

Reads were aligned to the GRCm39 mouse reference genome. Cell-by-gene count matrices were generated using the 10X Genomics CellRanger pipelines with default parameters. Data was then processed using CellBender^89^ with default parameters to remove background noise and the DoubletFinder R package^90^ to predict and remove doublets. Nuclei containing a minimum of 500 genes and less than 10% mitochondrial gene content were retained for analysis. Normalization and scaling were performed using the standard Seurat v5 pipeline in R, followed by PCA using the top 2000 variable genes. Cell type classification was performed using annotated P11 data^28^ as a reference object and a predicted threshold of 0.75. Clusters were visualized in 2D space using Uniform Manifold Approximation and Projection (UMAP).

### RNA data analysis

V1 data was obtained from: https://allen-developmental-mouse-atlas.s3.us-west-2.amazonaws.com/Multiome/DevVIS_multiome_snRNA_processed.h5ad^28^.

The data was subset using the *Pvalb Gaba* and *Sst Gaba* subclass labels at the following timepoints: P2, P5, P8, P11, P14, and P56.

#### Maturation analysis

To quantify transcriptional maturation, pseudobulk gene expression profiles were computed for each cell type (PV and SST) and developmental timepoint (P2, 5, 8, 11, 14) by averaging normalized expression values across cells. To reduce noise from lowly expressed genes, analyses were restricted to genes with average expression > 0.1 in at least 10% of cell type– timepoint groups (∼10,000 genes). This filtering reduced the influence of lowly expressed genes while preserving broad gene coverage for downstream analyses.

Maturation was assessed relative to adult (P56) reference profiles using two complementary metrics: Spearman correlation and Euclidean distance. Spearman correlation was used to quantify similarity in gene expression hierarchy (i.e., relative ranking of genes), while Euclidean distance measured differences in expression magnitude across genes. This procedure was repeated across 50 bootstrap iterations for each condition, and the mean and standard deviation of the resulting correlations were used to estimate maturation trajectories and variability. This approach controls for differences in cell number across conditions and provides a robust estimate of transcriptional similarity.

#### SCENIC GRN analysis

Gene regulatory network (GRN) analysis was performed using the SCENIC workflow implemented with pySCENIC^91^. Regulatory networks were inferred from snRNA-seq data using the GRNBoost2 algorithm to identify transcription factor (TF)-target co-expression modules.

Regulon pruning based on motif enrichment was subsequently performed using pySCENIC cisTarget databases, and regulon activity scores were calculated for each cell using AUCell. Regulons with normalized enrichment score (NES) values greater than 2.0 were retained for downstream analyses.

For subtype-specific analysis, PV INs were subset from the integrated Seurat object based on subclass_label annotations. Cells were grouped by developmental age (age_label) and differential regulon activity between P11 and P8 cells was assessed using the Wilcoxon rank-sum test implemented in Seurat (FindMarkers, assay = “pyscenic”). Regulon activity matrices were extracted from AUCell scores stored in the “pyscenic” assay.

To quantify effect size, mean regulon activity (AUC score) was computed separately for P11 and P8 cells for each TF regulon, and differential activity was defined as the difference in AUC between P11 and P8. Regulons were considered significantly differentially active if they satisfied both an adjusted *p*-value threshold of < 0.05 and an absolute ΔAUC > 0.02. Differentially active regulons were separated into upregulated and downregulated groups based on the sign of ΔAUC.

#### Pathway enrichment analysis

We computed gene-wise differences in expression magnitude and rank between developmental timepoints and the adult reference. Genes (non-zero average expression in at least 10% of cell type–timepoint groups) were classified based on their contribution to changes in expression magnitude (Euclidean), gene expression hierarchy (Spearman), or both. Specifically, genes with the largest changes in absolute expression difference were assigned to the magnitude category, whereas genes with the largest changes in rank difference were assigned to the gene program category; genes meeting both criteria were classified as contributing to both. Thresholds for classification were defined using percentile-based cutoffs (top 90%) within each metric.

Directionality of change was determined based on expression differences between developmental timepoints (e.g., P8 vs. P11), allowing genes to be further categorized as upregulated or downregulated during maturation. These gene sets were subsequently used for pathway enrichment analysis.

Pathway enrichment analysis was performed using over-representation analysis implemented in clusterProfiler^92^. Gene sets were obtained from MSigDB using the msigdb package (v26.1.0), restricting to curated Reactome pathways (C2:CP:REACTOME) for *Mus musculus*. For each gene category (e.g., magnitude-driven, gene program-driven, or both), enrichment was computed using the enricher function with a background defined as all genes passing the expression threshold used in the analysis. P-values were adjusted for multiple testing using the Benjamini–Hochberg procedure, and pathways with adjusted p-values below 0.05 were considered significant. For visualization, enrichment significance is reported as −log_10_(adjusted p-value), and gene counts indicate the number of genes contributing to each enriched pathway.

#### Gene bias analysis

Gene bias analysis was performed using pseudobulk expression profiles from the V1 dataset. For each cell type (PV and SST) and developmental timepoint, pseudobulk expression was computed by averaging normalized gene expression values across cells. Developmental changes in gene expression were quantified for each gene as the difference between timepoints (Δ expression), defined relative to the maturation window of interest (P11 – P8). To assess cell-type-specific bias, we computed the difference in developmental change between PV and SST interneurons (Δ effect size = Δ PV - Δ SST), where positive values indicate greater upregulation in PV and negative values indicate greater upregulation in SST.

Gene bias was evaluated with pathway-defined gene sets identified from enrichment analysis. For each pathway, genes were ranked by the magnitude of their bias, and the top-ranking genes were selected for visualization and functional categorization. Using the same gene sets and gene bias metric defined above, we performed an equivalent analysis in an independent S1 dataset spanning the P9-P11 developmental window.

### Immunohistochemistry

Mice were anesthetized with 5% isoflurane and transcardially perfused with ice cold phosphate buffer saline PBS (0.1 M) followed by ice cold 4% paraformaldehyde in PBS and post-fixed overnight at 4°C. Coronal sections (40 μm) were cut on a vibratome (Leica VT1200S). The following primary antibodies were used: mouse anti-parvalbumin (1:1,000, SAB4200545, Sigma Aldrich). Sections were imaged on an ApoTome2 microscope (Zen2 software, Zeiss; 25X objective, 0.3 NA) and analyzed with Fiji.

#### PV cell density quantification with Fiji

The primary somatosensory cortex was selected according to *Paxinos and Franklin’s the Mouse Brain in Stereotaxic Coordinates* (5th edition), and the area of the selected region was recorded. Images underwent thresholding to retain the top 3.2% brightest pixels and remove background. Bright neurites were manually removed to retain only PV-positive cell bodies. Images were then converted to binary masks and touching cell bodies were separated using the “Watershed” function. PV cell bodies were counted using the “Analyze Particles” function, with particle area set from 5 µm^2^ - infinity and circularity set from 0.20 - 1.00. PV cell density was calculated as the ratio of the number of PV-positive cells to the area of the region selected. For each mouse, the ipsilateral-to-contralateral ratio was calculated from the mean densities. Groups were compared using a two-sided Mann–Whitney U test because the distribution of mouse-level density ratios deviated from normality.

### Computational model

#### Network model and dynamics

We considered a firing-rate network model comprising three neuronal populations: 400 excitatory (Exc), 40 parvalbumin-positive inhibitory (PV), and 40 somatostatin-positive inhibitory (SST) neurons. Each population consisted of multiple neurons, each represented by a single firing-rate unit. We denote neuronal populations by Greek indices *α*, *β* ∈ {Exc, PV, SST} and neurons within each population by Latin indices *i* and *j*. The network dynamics followed a first-order rate model with rectified-linear transfer functions.

For neuron *i* in population *α*, the firing rate 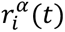 evolved according to

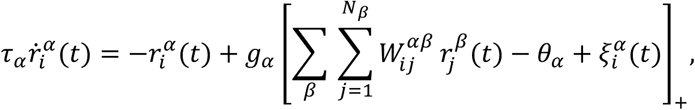

where *τ_α_* is the effective time constant of population *α*, *g_α_* is the population-specific gain, 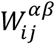 denotes the synaptic coupling from presynaptic neuron *j* in population *β* to postsynaptic neuron *i* in population *α*, *θ_α_* is the firing threshold, and [⋅]_+_ = max(0,⋅) denotes a rectified linear (ReLU) nonlinearity. The term 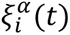 represents external fluctuations modeled as an Ornstein-Uhlenbeck (OU) process.

The synaptic connectivity is specified by the matrices *W^αβ^*, whose entries are the synaptic weights 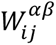. The full connectivity matrix is therefore written in block form as

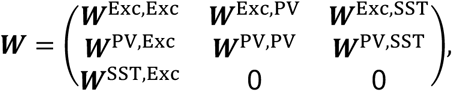

where each block ***W****^αβ^* has dimensions *N_α_* × *N_β_*, with *N_α_* and *N_β_* denoting the number of neurons in population *α* and *β*, respectively. We set to zero synaptic connections between cell types in populations that are negligible in the adult mouse^55–57^ assuming they are also negligible at P9-14.

#### Parameterization of the connectivity matrix

The network connectivity was parameterized by a population-level matrix of mean synaptic strengths: each block ***W****^αβ^* was initially constructed as a fully connected matrix whose entries were all equal to the corresponding mean value,

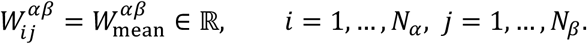

To introduce structured recurrent excitation, clustering was applied exclusively to the excitatory-to-excitatory block ***W***^Exc,Exc^. We defined *M* clusters, where *M* is a model parameter. Each excitatory neuron was assigned to exactly *K* clusters, with *K* fixed across neurons and treated as an independent model parameter. Denoting by 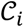 the set of clusters to which neuron *i* belongs, the excitatory synaptic weights were modified according to

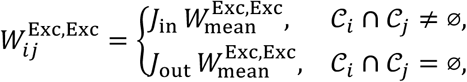

where *J*_in_ > 1 and *J*_out_ < 1 respectively strengthen and weaken recurrent excitatory connections^43^. The values of *J*_in_ and *J*_out_ were treated as independent model parameters.

Finally, we introduced heterogeneous connectivity by independently perturbing every synaptic weight. Specifically,

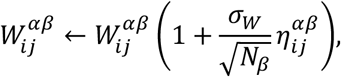

where 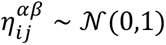 are independent Gaussian random variables and *σ*_W_ controls the amplitude of the heterogeneity and was treated as an independent parameter. The perturbation scales as 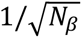 with the size of the presynaptic population, ensuring that the variance of the total synaptic input remains approximately independent of population size. The parameter *σ*_W_ was varied systematically in the analyses below.

#### Stochastic input model

We modeled external fluctuations as an Ornstein–Uhlenbeck (OU) process acting on each neuron. Defining

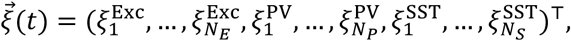

the stochastic input evolved according to:

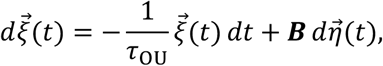

where 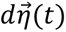 is a vector of independent Wiener increments. To determine the relationship between the noise amplitude ***B*** and the stationary covariance 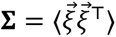, we computed the dynamics of the second moment. This yields:

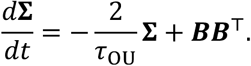

At stationarity, the covariance satisfies:

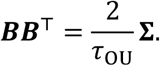

The covariance matrix can be factorized as **Σ** = ***LL***^T^ (e.g., via Cholesky decomposition), which allows us to write:

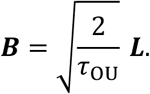

We computed ***L*** using *np.linalg.cholesky* and simulated the OU process via Euler–Maruyama:

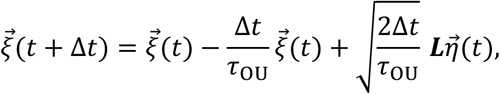

with 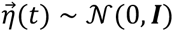.

#### Parameterization of the noise covariance

The stationary covariance matrix **Σ** was parameterized separately for each neuronal population, while correlations between different populations were neglected. Accordingly, **Σ** was block diagonal,

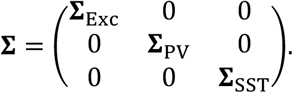

Excitatory neurons were divided into *M* equally sized clusters. Denoting by *c_i_* ∈ {1, … , *M*} the cluster assignment of neuron *i*, the covariance between two excitatory neurons depended only on the distance between their clusters. Specifically,

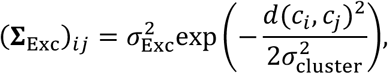

where:

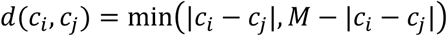

The parameter *σ*_cluster_ controls the correlation length across excitatory clusters, while for PV and SST populations, the covariance matrices were given by:

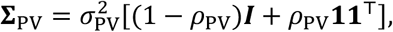

and

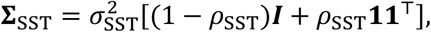

where *σ*_Exc_, *σ*_PV_, and *σ*_SST_ determine the marginal standard deviation of the fluctuations in each population, while *ρ*_PV_ and *ρ*_SST_ denote the pairwise correlation coefficients within the inhibitory populations.

#### Numerical simulations

The network dynamics were integrated using the forward Euler method with time step Δ*t* = 0.5 *s* ms.

Simulations were run for 30 s to capture spontaneous transitions between metastable states. Unless otherwise stated, all neurons were initialized with zero firing rate, ***r***(0) = **0**.

To quantify correlated activity, we computed the PCC between the firing-rate traces of all pairs of excitatory neurons and averaged the resulting coefficients.

To characterize the collective network dynamics, we defined the population-averaged excitatory firing rate,

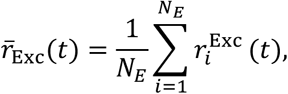

which was used throughout the analyses to quantify transitions between low- and high-activity states.

#### Quantification of bimodality of the average firing rate

We assessed whether the simulated neural trace, averaged over neurons, exhibited bistable dynamics characterized by transitions between low- and high-activity states. We identified local maxima of the estimated firing rate density and selected the two highest-density peaks as candidate DOWN- and UP-state firing rates. We required these peaks to be separated by a local minimum in the density estimate, which defined the threshold between the two states. The strength of bimodality was quantified by comparing the density at this minimum (*ρ*_valley_) with the mean density at the two peaks (*ρ*_peak,1_ and *ρ*_peak,2_),

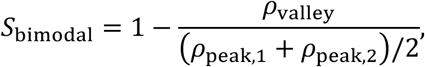

which ranges from 0 (poorly separated modes) to 1 (complete separation). Each time point was assigned to the DOWN or UP state according to whether its firing rate was below or above the estimated threshold. To assess whether the low-activity state was consistent with quiescence, we computed the mean, *μ*_down_, and standard deviation, *σ*_down_, of the firing rate within the DOWN state. We thus defined a continuous score describing the proximity of the DOWN-state firing rate to zero as:

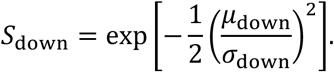

The overall bistability score was computed as the geometric mean of the bimodality and DOWN-state scores:

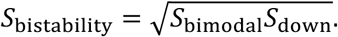

#### Parameter exploration and model selection

Model parameters were sampled independently from the uniform prior distributions listed in Table 1. In total, 10^5^ parameter sets were generated and simulated independently. For each parameter set, we computed summary statistics characterizing the network dynamics: the bistability score, the mean pairwise Pearson correlation coefficient (PCC), the frequency of SNEs, and the participation ratio of the excitatory population activity.

**TABLE 1:** Parameters used in simulations.

| Parameter | Uniform prior |
| --- | --- |
| $W_{\text{mean}}^{\text{Exc,Exc}}$ | $\mathcal{U}(1.0, 2.0)$ |
| $W_{\text{mean}}^{\text{Exc,PV}}$ | $\mathcal{U}(0.3, 1.5)$ |
| $W_{\text{mean}}^{\text{Exc,SST}}$ | $\mathcal{U}(0.1, 0.9)$ |
| $W_{\text{mean}}^{\text{PV,Exc}}$ | $\mathcal{U}(1.0, 1.6)$ |
| $W_{\text{mean}}^{\text{PV,PV}}$ | $\mathcal{U}(0.1, 0.5)$ |
| $W_{\text{mean}}^{\text{PV,SST}}$ | $\mathcal{U}(0.1, 0.5)$ |
| $W_{\text{mean}}^{\text{SST,Exc}}$ | $\mathcal{U}(0.1, 0.6)$ |
| $\theta_{\text{Exc}}$ | $\mathcal{U}(4, 8)$ |
| $\theta_{\text{PV}}$ | $\mathcal{U}(3, 12)$ |
| $\theta_{\text{SST}}$ | $\mathcal{U}(3, 9)$ |
| $\tau_{\text{Exc}}$ | $\mathcal{U}(10, 30)$ ms |
| $\tau_{\text{PV}}$ | $\mathcal{U}(2, 10)$ ms |
| $\tau_{\text{SST}}$ | $\mathcal{U}(10, 40)$ ms |
| $\tau_{\text{OU}}$ | $\mathcal{U}(1, 10)$ ms |
| $M$ | $\mathcal{U}_{\text{int}}(5, 15)$ |
| $K$ | $\mathcal{U}_{\text{int}}(1, 5)$ |
| $J_{\text{in}}$ | $\mathcal{U}(1.1, 3.5)$ |
| $J_{\text{out}}$ | $\mathcal{U}(0.2, 0.9)$ |
| $\sigma_W$ | $\mathcal{U}(0, 10)$ |
| $\rho_{\text{PV}}$ | $\mathcal{U}(0, 0.8)$ |
| $\rho_{\text{SST}}$ | $\mathcal{U}(0, 0.8)$ |

To identify parameter sets that best reproduced the experimental observations, we defined a score quantifying the agreement between the simulated and experimentally measured summary statistics at postnatal days P9 and P14. Specifically, the score was the L2 distance between the statistics of the model’s output (PCC Exc-Exc, Participation ratio, frequency of SNE) and the statistics of the data. The 10^4^ best-fitting pairs of parameter sets (***θ***_P9_, ***θ***_P14_) were retained for further analysis. Among these candidate pairs, we quantified the biological plausibility of the developmental parameter changes. Let

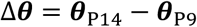

denote the parameter change between developmental stages. Parameters expected to increase during development, namely the synaptic projections from Exc and PV neurons^58,59^ and the Exc and PV thresholds^60^, were assigned to the set ℐ. Conversely, parameters expected to remain approximately constant, namely the synaptic projections from SST neurons and the SST threshold^61^, were assigned to the set S (Table 1). The other parameters were left unconstrained. We then defined the biological alignment score

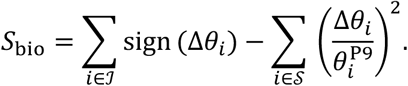

The first term rewards developmental changes occurring in the expected direction, while the second penalizes relative changes in parameters expected to remain stable. Candidate model pairs were ranked according to *S*_bio_, and the 500 highest-scoring pairs were retained. For each of the 500 selected model pairs, we reconstructed the developmental trajectory of the model parameters by assuming linear changes between P9 and P14. Specifically, for each parameter *θ_i_*, intermediate values were obtained by linear interpolation,

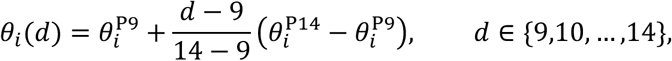

where *d* denotes the postnatal day. The trajectory may be non-linear, so that an even better fit with the data may emerge. This procedure yielded a complete developmental trajectory for every parameter in each selected model.

The network was then simulated independently at each developmental stage (P9-P14) using the corresponding interpolated parameter values. The same simulation and analysis pipeline described above was applied to compute the bistability score, mean pairwise Pearson correlation coefficient (PCC), frequency of SNEs, and participation ratio of the excitatory population activity throughout development.

### Optogenetic perturbations in the model

We simulated optogenetic perturbations by adding a constant external input to all neurons within a selected population. The dynamics of neuron *i* in population *α* became:

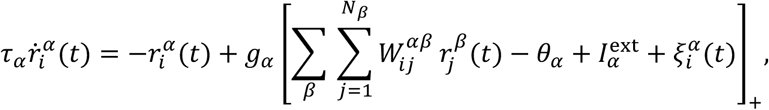

where 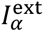 denotes the optogenetic input delivered uniformly to all neurons in population *α*. Equivalently, this perturbation can be interpreted as a shift of the effective firing threshold.

We considered three stimulation conditions: no stimulation (control), selective activation of PV neurons, and selective activation of SST neurons. The stimulation amplitude was varied between 0 and 1.5. To quantify the effect of optogenetic stimulation on network dynamics, transitions between low- and high-activity states were identified using the same threshold determined from the corresponding control simulation 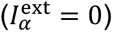.

## Notes

### Competing Interest Statement

The authors have declared no competing interest.

## REFERENCES

1. Leighton, A. H. & Lohmann, C. The Wiring of Developing Sensory Circuits—From Patterned Spontaneous Activity to Synaptic Plasticity Mechanisms. Front. Neural Circuits 10, (2016).

2. Luhmann, H. J. & Khazipov, R. Neuronal Activity Patterns in the Developing Barrel Cortex. Neuroscience 368, 256–267 (2018).

3. Cossart, R. & Garel, S. Step by step: cells with multiple functions in cortical circuit assembly. Nat. Rev. Neurosci. 23, 395–410 (2022).

4. Wu, M. W., Kourdougli, N. & Portera-Cailliau, C. Network state transitions during cortical development. Nat. Rev. Neurosci. 25, 535–552 (2024).

5. Golshani, P. et al. Internally Mediated Developmental Desynchronization of Neocortical Network Activity. J. Neurosci. 29, 10890–10899 (2009).

6. Rochefort, N. L. et al. Sparsification of neuronal activity in the visual cortex at eye-opening. Proc. Natl. Acad. Sci. 106, 15049–15054 (2009).

7. Siegel, F., Heimel, J. A., Peters, J. & Lohmann, C. Peripheral and Central Inputs Shape Network Dynamics in the Developing Visual Cortex In Vivo. Curr. Biol. 22, 253–258 (2012).

8. Che, A. et al. Layer I Interneurons Sharpen Sensory Maps during Neonatal Development. Neuron 99, 98–116.e7 (2018).

9. Nakazawa, S., Yoshimura, Y., Takagi, M., Mizuno, H. & Iwasato, T. Developmental Phase Transitions in Spatial Organization of Spontaneous Activity in Postnatal Barrel Cortex Layer 4. J. Neurosci. 40, 7637–7650 (2020).

10. Gjorgjieva, J., Mease, R. A., Moody, W. J. & Fairhall, A. L. Intrinsic Neuronal Properties Switch the Mode of Information Transmission in Networks. PLOS Comput. Biol. 10, e1003962 (2014).

11. Lakhera, S., Herbert, E. & Gjorgjieva, J. Modeling the Emergence of Circuit Organization and Function during Development. Cold Spring Harb. Perspect. Biol. 17, a041511 (2025).

12. Hanganu, I. L., Ben-Ari, Y. & Khazipov, R. Retinal Waves Trigger Spindle Bursts in the Neonatal Rat Visual Cortex. J. Neurosci. 26, 6728–6736 (2006).

13. Colonnese, M. T. et al. A Conserved Switch in Sensory Processing Prepares Developing Neocortex for Vision. Neuron 67, 480–498 (2010).

14. Ackman, J. B., Burbridge, T. J. & Crair, M. C. Retinal waves coordinate patterned activity throughout the developing visual system. Nature 490, 219–225 (2012).

15. Murata, Y. & Colonnese, M. T. An excitatory cortical feedback loop gates retinal wave transmission in rodent thalamus. eLife 5, e18816 (2016).

16. Field, D. J. What Is the Goal of Sensory Coding? Neural Comput. 6, 559–601 (1994).

17. Olshausen, B. A. & Field, D. J. Emergence of simple-cell receptive field properties by learning a sparse code for natural images. Nature 381, 607–609 (1996).

18. Olshausen, B. A. & Field, D. J. Sparse coding of sensory inputs. Curr. Opin. Neurobiol. 14, 481–487 (2004).

19. Babadi, B. & Sompolinsky, H. Sparseness and Expansion in Sensory Representations. Neuron 83, 1213–1226 (2014).

20. Stringer, C., Pachitariu, M., Steinmetz, N., Carandini, M. & Harris, K. D. High-dimensional geometry of population responses in visual cortex. Nature 571, 361–365 (2019).

21. Stringer, C. et al. Spontaneous behaviors drive multidimensional, brainwide activity. Science 364, eaav7893 (2019).

22. Majnik, J. et al. Longitudinal tracking of neuronal activity from the same cells in the developing brain using Track2p. eLife 14, (2025).

23. Wamsley, B. & Fishell, G. Genetic and activity-dependent mechanisms underlying interneuron diversity. Nat. Rev. Neurosci. 18, 299–309 (2017).

24. Pouchelon, G. et al. The organization and development of cortical interneuron presynaptic circuits are area specific. Cell Rep. 37, 109993 (2021).

25. García, N. V. D. M. & Fishell, G. Interneuron Diversity: How Form Becomes Function. Cold Spring Harb. Perspect. Biol. 17, a041513 (2025).

26. Wonders, C. P. & Anderson, S. A. The origin and specification of cortical interneurons. Nat. Rev. Neurosci. 7, 687–696 (2006).

27. Lim, L., Mi, D., Llorca, A. & Marín, O. Development and Functional Diversification of Cortical Interneurons. Neuron 100, 294–313 (2018).

28. Gao, Y. et al. Continuous cell-type diversification in mouse visual cortex development. Nature 647, 127–142 (2025).

29. Favuzzi, E. et al. Distinct molecular programs regulate synapse specificity in cortical inhibitory circuits. Science 363, 413–417 (2019).

30. Chini, M., Pfeffer, T. & Hanganu-Opatz, I. An increase of inhibition drives the developmental decorrelation of neural activity. eLife 11, e78811 (2022).

31. Mòdol, L., Moissidis, M., Selten, M., Oozeer, F. & Marín, O. Somatostatin interneurons control the timing of developmental desynchronization in cortical networks. Neuron 0, (2024).

32. Wosniack, M. E. et al. Adaptation of spontaneous activity in the developing visual cortex. eLife 10, e61619 (2021).

33. Dwulet, J. M. et al. Correlated spontaneous activity sets up multi-sensory integration in the developing higher-order cortex. 2024.07.19.603239 Preprint at 10.1101/2024.07.19.603239 (2026).

34. Teh, K. L., Sibille, J., Gehr, C. & Kremkow, J. Retinal waves align the concentric orientation map in mouse superior colliculus to the center of vision. Sci. Adv. 9, eadf4240 (2023).

35. Rahmati, V., Kirmse, K., Holthoff, K., Schwabe, L. & Kiebel, S. J. Developmental Emergence of Sparse Coding: A Dynamic Systems Approach. Sci. Rep. 7, 13015 (2017).

36. Romagnoni, A., Colonnese, M. T., Touboul, J. D. & Gutkin, B. S. Progressive alignment of inhibitory and excitatory delay may drive a rapid developmental switch in cortical network dynamics. J. Neurophysiol. 123, 1583–1599 (2020).

37. Ko, H. et al. The emergence of functional microcircuits in visual cortex. Nature 496, 96– 100 (2013).

38. O’Donnell, C., Gonçalves, J. T., Portera-Cailliau, C. & Sejnowski, T. J. Beyond excitation/inhibition imbalance in multidimensional models of neural circuit changes in brain disorders. eLife 6, e26724 (2017).

39. Tahvili, F., Vinck, M. & Volo, M. di. PV and SOM cells play distinct causal roles in controlling network oscillations and stability. Cell Rep. 44, (2025).

40. Parker, M. M., Rubin, J. E. & Huang, C. State modulation in spatial networks with three interneuron subtypes. Sci. Adv. 11, eads9134 (2025).

41. Romero-Sosa, J. L., Motanis, H. & Buonomano, D. V. Differential Excitability of PV and SST Neurons Results in Distinct Functional Roles in Inhibition Stabilization of Up States. J. Neurosci. 41, 7182–7196 (2021).

42. Jercog, D. et al. UP-DOWN cortical dynamics reflect state transitions in a bistable network. eLife 6, e22425 (2017).

43. Litwin-Kumar, A. & Doiron, B. Slow dynamics and high variability in balanced cortical networks with clustered connections. Nat. Neurosci. 15, 1498–1505 (2012).

44. Shimamura, K., Hartigan, D. J., Martinez, S., Puelles, L. & Rubenstein, J. L. R. Longitudinal organization of the anterior neural plate and neural tube. Development 121, 3923–3933 (1995).

45. Modol, L. et al. Assemblies of Perisomatic GABAergic Neurons in the Developing Barrel Cortex. Neuron 105, 93–105.e4 (2020).

46. Recanatesi, S., Bradde, S., Balasubramanian, V., Steinmetz, N. A. & Shea-Brown, E. A scale-dependent measure of system dimensionality. Patterns 3, 100555 (2022).

47. Kourdougli, N. et al. Improvement of sensory deficits in fragile X mice by increasing cortical interneuron activity after the critical period. Neuron 0, (2023).

48. Bollmann, Y. et al. Prominent in vivo influence of single interneurons in the developing barrel cortex. Nat. Neurosci. 26, 1555–1565 (2023).

49. de la Rocha, J., Doiron, B., Shea-Brown, E., Josić, K. & Reyes, A. Correlation between neural spike trains increases with firing rate. Nature 448, 802–806 (2007).

50. Juarez, P., Salcedo-Arellano, M. J., Dufour, B. & Martinez-Cerdeño, V. Fragile X cortex is characterized by decreased parvalbumin-expressing interneurons. Cereb. Cortex N. Y. NY 34, bhae103 (2024).

51. Contractor, A., Ethell, I. M. & Portera-Cailliau, C. Cortical interneurons in autism. Nat. Neurosci. 24, 1648–1659 (2021).

52. Selby, L., Zhang, C. & Sun, Q.-Q. Major Defects in Neocortical GABAergic Inhibitory Circuits in Mice Lacking the Fragile X Mental Retardation Protein. Neurosci. Lett. 412, 227–232 (2007).

53. Wen, T. H. et al. Genetic Reduction of Matrix Metalloproteinase-9 Promotes Formation of Perineuronal Nets Around Parvalbumin-Expressing Interneurons and Normalizes Auditory Cortex Responses in Developing Fmr1 Knock-Out Mice. Cereb. Cortex N. Y. NY 28, 3951– 3964 (2018).

54. Filice, F., Vörckel, K. J., Sungur, A. Ö., Wöhr, M. & Schwaller, B. Reduction in parvalbumin expression not loss of the parvalbumin-expressing GABA interneuron subpopulation in genetic parvalbumin and shank mouse models of autism. Mol. Brain 9, 10 (2016).

55. Campagnola, L. et al. Local connectivity and synaptic dynamics in mouse and human neocortex. Science 375, eabj5861 (2022).

56. Billeh, Y. N. et al. Systematic Integration of Structural and Functional Data into Multi-scale Models of Mouse Primary Visual Cortex. Neuron 106, 388–403.e18 (2020).

57. Pfeffer, C. K., Xue, M., He, M., Huang, Z. J. & Scanziani, M. Inhibition of inhibition in visual cortex: the logic of connections between molecularly distinct interneurons. Nat. Neurosci. 16, 1068–1076 (2013).

58. Lecea, L. de, del Río, J. & Soriano, E. Developmental expression of parvalbumin mRNA in the cerebral cortex and hippocampus of the rat. Mol. Brain Res. 32, 1–13 (1995).

59. Maravall, M., Stern, E. A. & Svoboda, K. Development of intrinsic properties and excitability of layer 2/3 pyramidal neurons during a critical period for sensory maps in rat barrel cortex. J. Neurophysiol. 92, 144–156 (2004).

60. Okaty, B. W., Miller, M. N., Sugino, K., Hempel, C. M. & Nelson, S. B. Transcriptional and Electrophysiological Maturation of Neocortical Fast-Spiking GABAergic Interneurons. J. Neurosci. 29, 7040–7052 (2009).

61. Tuncdemir, S. N. et al. Early Somatostatin Interneuron Connectivity Mediates the Maturation of Deep Layer Cortical Circuits. Neuron 89, 521–535 (2016).

62. Mazzucato, L., Fontanini, A. & Camera, G. L. Dynamics of Multistable States during Ongoing and Evoked Cortical Activity. J. Neurosci. 35, 8214–8231 (2015).

63. Dipoppa, M. et al. Vision and Locomotion Shape the Interactions between Neuron Types in Mouse Visual Cortex. Neuron 98, 602–615.e8 (2018).

64. Keller, A. J. et al. A Disinhibitory Circuit for Contextual Modulation in Primary Visual Cortex. Neuron 108, 1181–1193.e8 (2020).

65. Martini, F. J., Guillamón-Vivancos, T., Moreno-Juan, V., Valdeolmillos, M. & López-Bendito, G. Spontaneous activity in developing thalamic and cortical sensory networks. Neuron 109, 2519–2534 (2021).

66. Ferrer, C. & De Marco García, N. V. The Role of Inhibitory Interneurons in Circuit Assembly and Refinement Across Sensory Cortices. Front. Neural Circuits 16, (2022).

67. Kroon, T., van Hugte, E., van Linge, L., Mansvelder, H. D. & Meredith, R. M. Early postnatal development of pyramidal neurons across layers of the mouse medial prefrontal cortex. Sci. Rep. 9, 5037 (2019).

68. Duan, Z. R. S. et al. GABAergic Restriction of Network Dynamics Regulates Interneuron Survival in the Developing Cortex. Neuron 105, 75–92.e5 (2020).

69. Picardo, M. A. et al. Pioneer GABA Cells Comprise a Subpopulation of Hub Neurons in the Developing Hippocampus. Neuron 71, 695–709 (2011).

70. Wang, C.-Z. et al. Early-generated interneurons regulate neuronal circuit formation during early postnatal development. eLife 8, e44649 (2019).

71. Nomura, T. et al. Delayed Maturation of Fast-Spiking Interneurons Is Rectified by Activation of the TrkB Receptor in the Mouse Model of Fragile X Syndrome. J. Neurosci. 37, 11298–11310 (2017).

72. Du, J., Zhang, L., Weiser, M., Rudy, B. & McBain, C. Developmental expression and functional characterization of the potassium-channel subunit Kv3.1b in parvalbumin-containing interneurons of the rat hippocampus. J. Neurosci. 16, 506–518 (1996).

73. Rudy, B. & McBain, C. J. Kv3 channels: voltage-gated K+ channels designed for high-frequency repetitive firing. Trends Neurosci. 24, 517–526 (2001).

74. Anastasiades, P. G. et al. GABAergic interneurons form transient layer-specific circuits in early postnatal neocortex. Nat. Commun. 7, 10584 (2016).

75. Freund, T. F. & Katona, I. Perisomatic Inhibition. Neuron 56, 33–42 (2007).

76. Hu, H., Gan, J. & Jonas, P. Fast-spiking, parvalbumin+ GABAergic interneurons: From cellular design to microcircuit function. Science 345, 1255263 (2014).

77. Silberberg, G. & Markram, H. Disynaptic Inhibition between Neocortical Pyramidal Cells Mediated by Martinotti Cells. Neuron 53, 735–746 (2007).

78. Urban-Ciecko, J. & Barth, A. L. Somatostatin-expressing neurons in cortical networks. Nat. Rev. Neurosci. 17, 401–409 (2016).

79. Gonçalves, J. T., Anstey, J. E., Golshani, P. & Portera-Cailliau, C. Circuit level defects in the developing neocortex of Fragile X mice. Nat. Neurosci. 16, 903–909 (2013).

80. Cheyne, J. E., Zabouri, N., Baddeley, D. & Lohmann, C. Spontaneous Activity Patterns Are Altered in the Developing Visual Cortex of the Fmr1 Knockout Mouse. Front. Neural Circuits 13, 57 (2019).

81. Renart, A. et al. The Asynchronous State in Cortical Circuits. Science 327, 587–590 (2010).

82. Rosenbaum, R., Smith, M. A., Kohn, A., Rubin, J. E. & Doiron, B. The spatial structure of correlated neuronal variability. Nat. Neurosci. 20, 107–114 (2017).

83. Engel, T. A. et al. Selective modulation of cortical state during spatial attention. Science 354, 1140–1144 (2016).

84. Shi, Y.-L., Steinmetz, N. A., Moore, T., Boahen, K. & Engel, T. A. Cortical state dynamics and selective attention define the spatial pattern of correlated variability in neocortex. Nat. Commun. 13, 44 (2022).

85. Cruz-Martin, A. & Portera-Cailliau, C. In Vivo Imaging of Axonal and Dendritic Structures in Neonatal Mouse Cortex. Cold Spring Harb. Protoc. 2014, pdb.prot080150 (2014).

86. Pachitariu, M. et al. Suite2p: beyond 10,000 neurons with standard two-photon microscopy. 061507 Preprint at 10.1101/061507 (2017).

87. Litwin-Kumar, A., Harris, K. D., Axel, R., Sompolinsky, H. & Abbott, L. F. Optimal Degrees of Synaptic Connectivity. Neuron 93, 1153–1164.e7 (2017).

88. Seabold, S. & Perktold, J. Statsmodels: Econometric and Statistical Modeling with Python. SciPy 2010 https://doi.org/10.25080/Majora-92bf1922-011 (2010) doi:10.25080/Majora-92bf1922-011.

89. Fleming, S. J. et al. Unsupervised removal of systematic background noise from droplet-based single-cell experiments using CellBender. Nat. Methods 20, 1323–1335 (2023).

90. McGinnis, C. S., Murrow, L. M. & Gartner, Z. J. DoubletFinder: Doublet Detection in Single-Cell RNA Sequencing Data Using Artificial Nearest Neighbors. Cell Syst. 8, 329–337.e4 (2019).

91. Van de Sande, B. et al. A scalable SCENIC workflow for single-cell gene regulatory network analysis. Nat. Protoc. 15, 2247–2276 (2020).

92. Xu, S. et al. Using clusterProfiler to characterize multiomics data. Nat. Protoc. 19, 3292– 3320 (2024).

93. Stringer, C. et al. Rastermap: a discovery method for neural population recordings. Nat. Neurosci. 28, 201–212 (2025).

94. Pospisil, D. A. & Pillow, J. W. Revisiting the high-dimensional geometry of population responses in the visual cortex. Proc. Natl. Acad. Sci. 122, e2506535122 (2025).

