## Supplemental Figures for "Parvalbumin interneuron maturation reorganizes cortical population activity structure during development"

**SUPPLEMENTARY FIGURES AND LEGENDS**

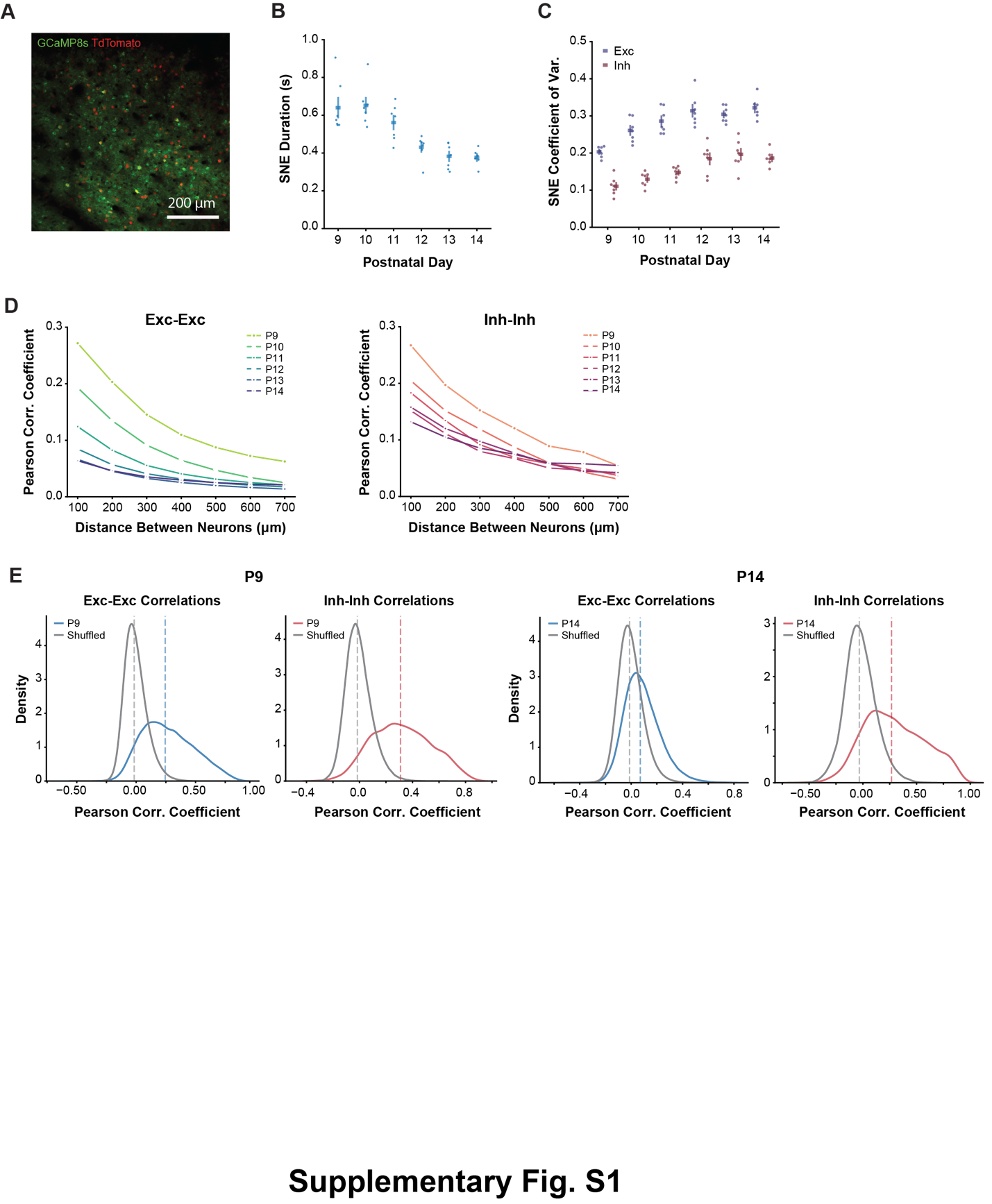

**Supplementary Fig. S1: Exc-Exc and Inh-Inh correlations are distance-dependent and higher than chance.**

(A) Example field of view (FOV) at P9. GCaMP8s expression in green, TdTomato expression in red (Nkx2.1+ MGE-derived interneurons).

(B) Synchronous network event (SNE) duration (seconds) decreases across the second postnatal week. LMM shows significant main effect of Day (Wald χ^2^ = 122.57, p = 8.96x10^-25^).

(C) Coefficient of variation (CV) increases for both Exc and Inh neurons. Significant main effects of Day (Wald χ^2^ = 151.93, p = 5.18x10^-31^), Cell Type (Wald χ^2^ = 62.91, p = 2.16x10^-15^), and a significant Day x Cell Type interaction (Wald χ^2^ = 11.84, p = 0.037). Post hoc pairwise comparisons between Exc and Inh within each postnatal day were significant for all days (adjusted p < 0.001).

(D) Pearson correlation coefficients (PCC) binned by distance between neuron pairs. Left: Exc-Exc. Distance-dependence decreases across the second postnatal week. Right: Inh-Inh. Distance-dependence was significant at all postnatal days for both Exc-Exc and Inh-Inh neuron pairs (Fisher's combined p < 0.001). The strength of distance-dependence decreased significantly with age for both Exc-Exc pairs (β = −0.074, p < 0.001) and Inh-Inh pairs (β = −0.048, p < 0.001), with Exc-Exc showing a significantly steeper decline (interaction β = −0.026, p = 0.021).

(E) Distribution of PCCs for Exc-Exc and Inh-Inh in experimental data vs. shuffled data (chance) at P9 (left) and P14 (right). Dashed lines indicate mean. Exc-Exc and Inh-Inh correlations are higher than would be expected by chance at P09 (Exc: p < 0.001, Inh: p < 0.001) and at P14 (Exc: p < 0.001, Inh: p = 0.018).

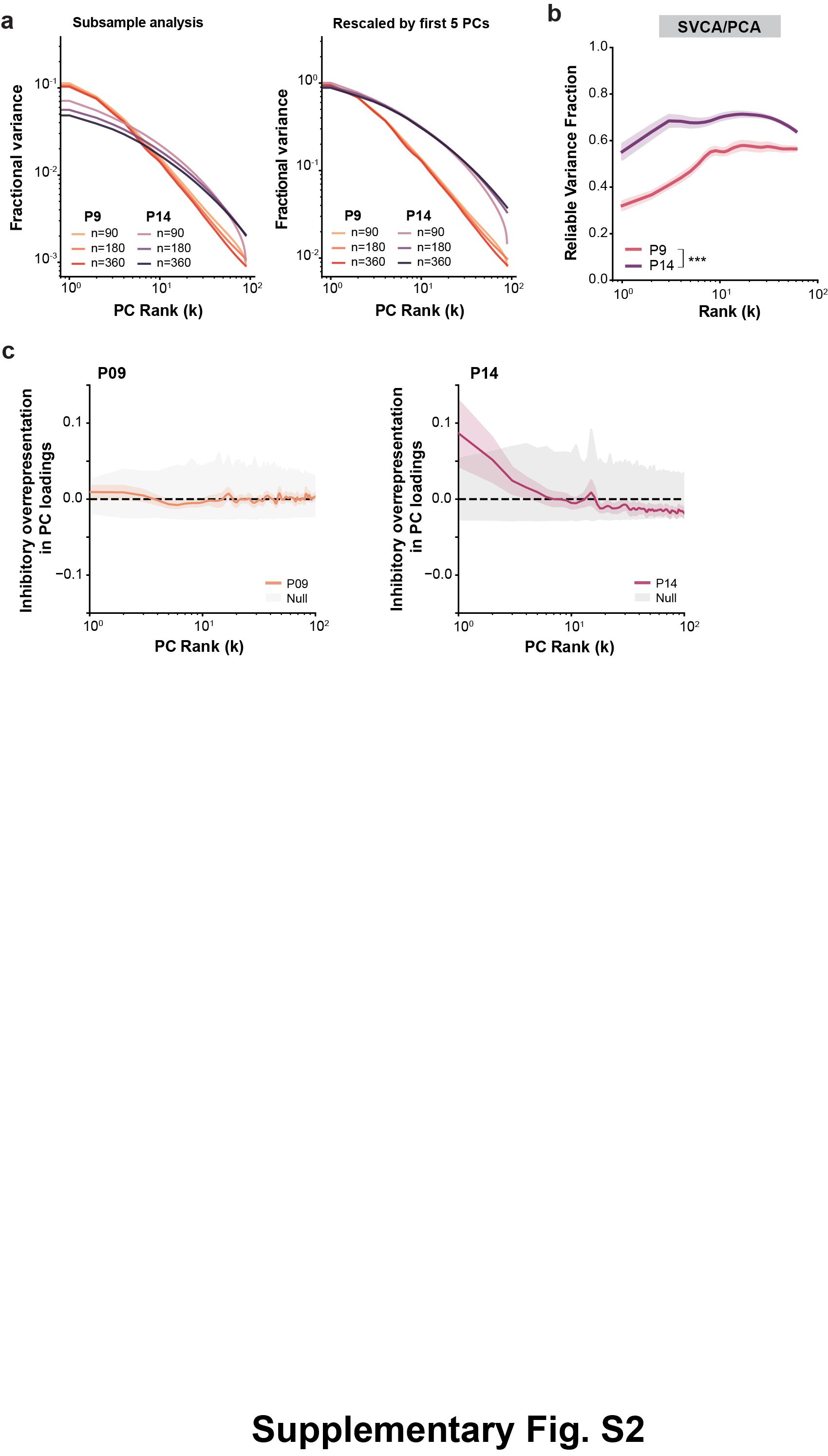

**Supplementary Fig. S2: Eigenspectra are stable across subsamples.**

(A) Fractional variance explained by descending ranks of principal components (PCs) for P9 and P14 using different subsample sizes: 90, 180, and 360 neurons (left). Fractional variance rescaled by the mean variance of the top 5 PCs (right). The different subsamples collapse along an intermediate range of PCs at P14. Mean curves shown. n=7 mice for P9-P13, n=6 mice for P14; 500-900 neurons per FOV.

(B) Reliable variance fraction (ratio of SVCA fractional shared variance to PCA fractional variance at matched ranks). LMM reveals significant main effect of Day at early and intermediate ranks (Early: β = 0.260, p < 0.001; Intermediate: β = 0.113, p < 0.001).

(C) Inhibitory overrepresentation in PC loadings as a function of PC rank at P9 (left) and P14 (right) with mean-centered activity (matching the pre-processing of eigenspectrum analyses), relative to a null distribution generated by randomly permuting cell type labels (gray). Color shaded regions indicate mean ± SEM. Dashed line indicates 0 (proportional representation).

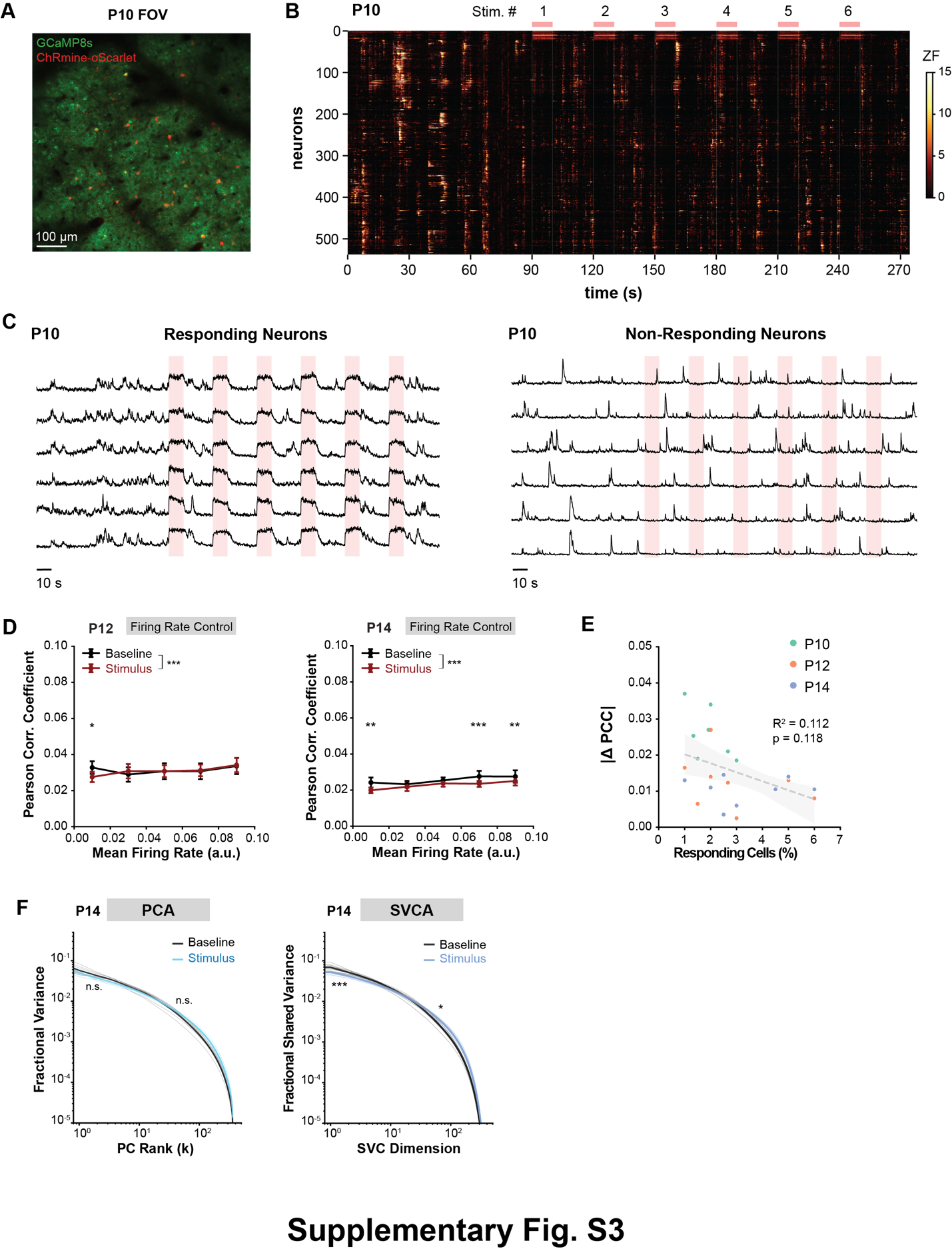

**Supplementary Fig. S3: PV interneuron activation validation.**

(A) Example FOV at P10 showing GCaMP8s (green) and ChRmine-oScarlet expression (red).

(B) Example raster of activity after optogenetic activation of PV INs at P10.

(C) Example traces from stimulus responsive putative PV INs (left) and stimulus non-responsive neurons (right); vertical red bars indicate epochs of optogenetic stimulation. Each trace was independently normalized to its maximum absolute response.

(D) Exc-Exc PCC binned by mean firing rate during baseline and stimulus periods at P12 (left) and P14 (right). Points represent the mean across mice, and error bars indicate SEM across mice. Correlations were computed from an equal number of randomly sampled neuron pairs per recording (see Methods). Significant interactions were followed by post hoc LMMs comparing Baseline and Stimulus within each firing-rate bin. At P12, significant effects were observed for Firing-Rate Bin (Wald χ^2^ = 20.90, p = 3.31x10^-4^), Condition (Wald χ^2^ = 14.67, p = 1.28x10^-4^), and the Firing-Rate Bin × Condition interaction (Wald χ^2^ = 18.69, p = 9.06 x10^-4^). Post hoc comparisons were significant for only the 0.01 firing-rate bin (adjusted p = 0.012). At P14, significant effects were observed for Firing-Rate Bin (Wald χ^2^ = 27.52, p = 1.56x10^-5^), Condition (Wald χ^2^ = 19.17, p = 1.20x10^-5^), and the Firing-Rate Bin × Condition interaction (Wald χ^2^ = 10.52, p = 0.033). Post hoc comparisons were significant for the following firing-rate bins (adjusted p < 0.005): 0.01, 0.07, 0.09.

(E) Relationship between number of responding PV INs (as a percent of the total population) and the change in PCC colored by timepoint. Linear regression provided no evidence that the percentage of responding cells predicted the effect on PCC (b = −0.002, 95% CI [−0.005, 0.001], t(21)=−1.63, p = 0.118, R^2^ = 0.112).

(F) Left, Fraction of variance explained by descending ranks of PCs for baseline vs. PV IN activation at P14 (left). No significant difference between baseline and PV IN activation (Early: β = -0.099, p = 0.095; Intermediate: β = 0.073, p = 0.204). Right, Fraction of reliable variance explained by descending ranks of PCs for baseline vs. PV IN activation at P14 (right). Significant difference between baseline and PV IN activation at early and intermediate ranks (Early: β = -0.189, p < 0.001; Intermediate: β = 0.102, p = 0.045).

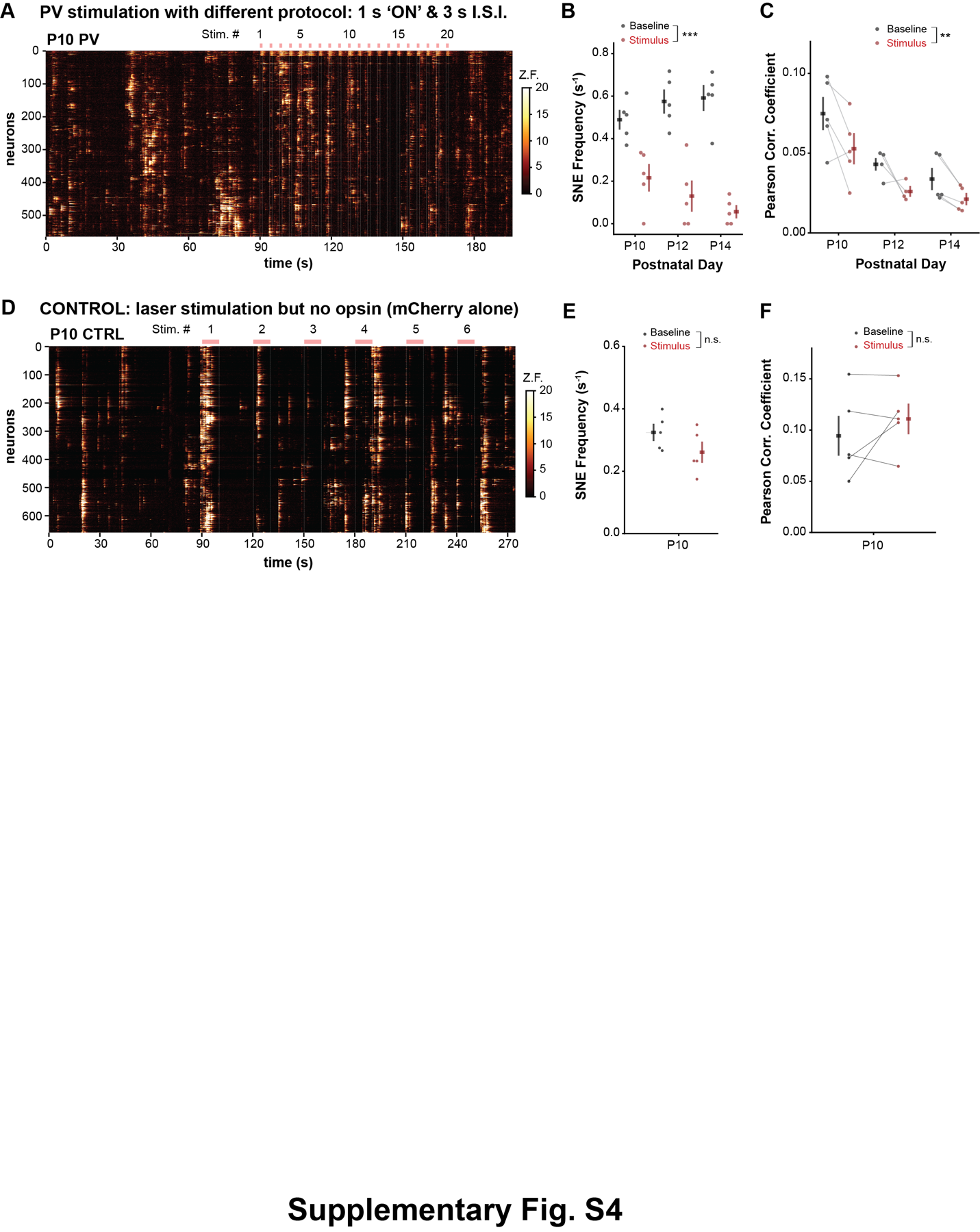

**Supplementary Fig. S4: Effects of PV interneuron activation are robust to protocol and are not due to laser artifact.**

(A) Example rasters with an alternative stimulation protocol (20 trials, 1s on / 3s off) at P10.

(B) SNE frequency during baseline vs. PV IN activation with alternative protocol. LMM shows significant main effects of Condition (Wald χ^2^ = 14.81, p = 1.19x10^-4^) but not Day (Wald χ^2^ = 1.62, p = 0.445), and a significant Day x Condition interaction (Wald χ^2^ = 6.20, p = 0.045). Post hoc contrasts showed significant effects of optogenetics at each day, increasing over time (P10: *p* = 1.19x10^-4^, P12: *p* = 5.50×10⁻^10^; P14: *p* = 1.05×10⁻^12^). n = 5 mice, 500-900 neurons per FOV.

(C) Exc-Exc PCC during baseline and during PV IN activation. Significant main effects of Day (Wald χ^2^ = 28.77, p = 5.65x10^-7^) and Condition (Wald χ^2^ = 6.83, p = 0.009).

(D) Example raster at P10 from a control experiment where mCherry was virally expressed alone (no opsin).

(E) SNE frequency during baseline and stimulation in control animals showed no significant effect of stimulation (p = 0.051). n = 5 mice, 500-900 neurons per FOV.

(F) PCC during baseline and stimulation in control animals showed no significant effect of stimulation (p = 0.362).

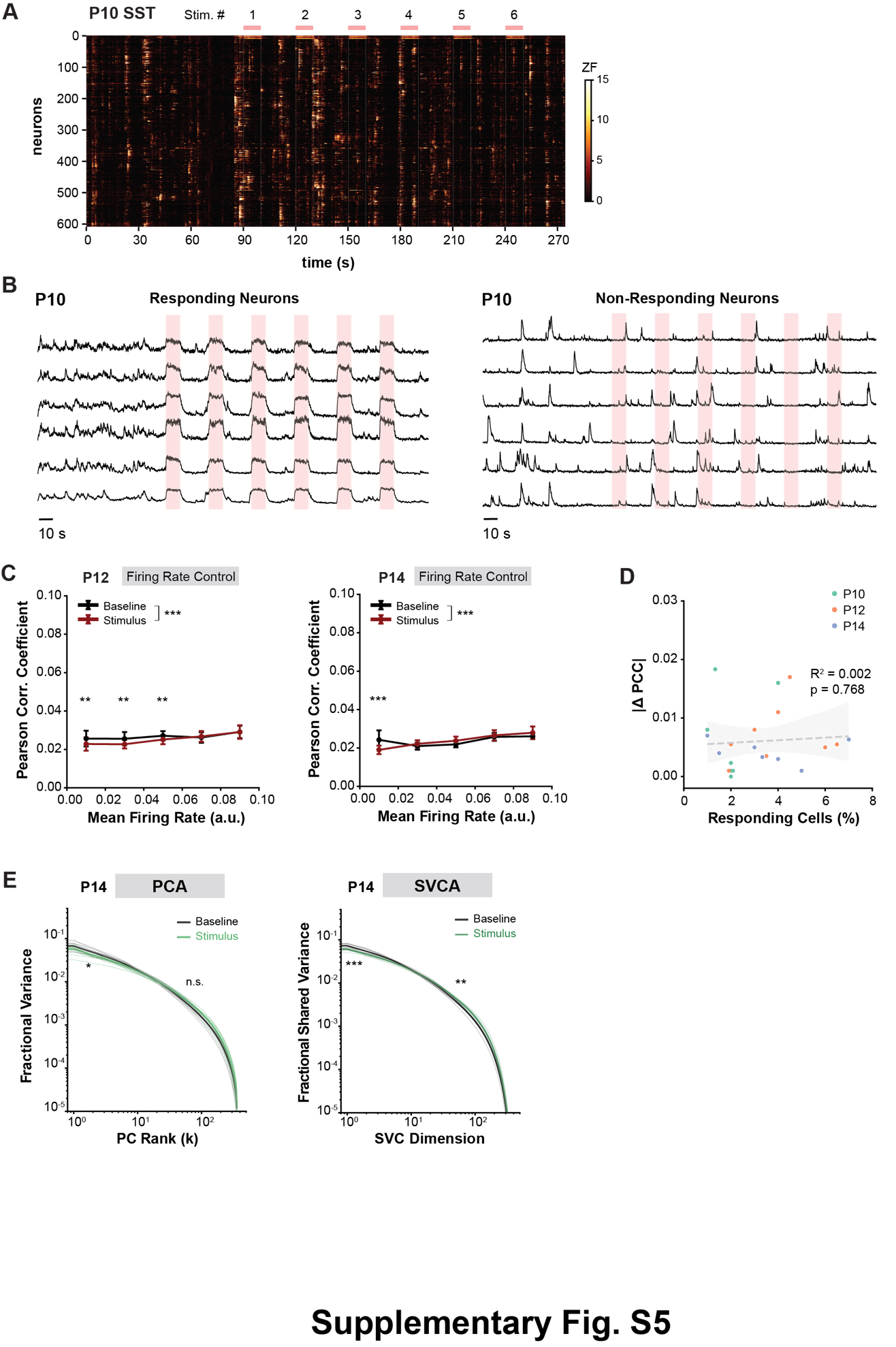

**Supplementary Fig. S5: SST interneuron activation validation.**

(A) Example raster of activity after optogenetic activation of SST INs at P10.

(B) Example traces from stimulus responsive putative SST INs (left) and stimulus non-responsive neurons (right). Each trace was independently normalized to its maximum absolute response.

(C) Exc-Exc PCC binned by mean firing rate during baseline and stimulus periods at P12 (left) and P14 (right). Points represent the mean across mice, and error bars indicate SEM across mice. Correlations were computed from an equal number of randomly sampled neuron pairs per recording and statistics were performed using LMMs. Significant interactions were followed by post hoc LMMs comparing Baseline and Stimulus within each firing-rate bin. At P12, significant effects were observed for Firing-Rate Bin (Wald χ^2^ = 22.99, p = 1.27 x10^-4^), Condition (Wald χ^2^ = 14.42, p = 1.46 x10^-4^), and the Firing-Rate Bin × Condition interaction (Wald χ^2^ = 12.38, p = 0.015). Post hoc comparisons were significant for the following firing-rate bins (adjusted p < 0.005): 0.01, 0.03, 0.05. At P14, significant effects were observed for Firing-Rate Bin (Wald χ^2^ = 43.17, p = 9.53 x10^-9^), Condition (Wald χ^2^ = 41.08, p = 1.46 x10^-10^), and the Firing-Rate Bin × Condition interaction (Wald χ^2^ = 46.58, p = 1.86 x10^-9^). Post comparisons were significant only for the 0.01 firing-rate bin (adjusted p < 0.001).

(D) Relationship between number of responding SST INs (percent of the total population) and the change in PCC colored by timepoint. Linear regression provided no evidence that the percentage of responding cells predicted the effect on PCC (b = −0.0002, 95% CI [−0.001, 0.001], p = 0.768, R^2^ = 0.002).

(E) Left, Fraction of variance explained by descending ranks of PCs for baseline vs. PV IN activation at P14 (left). LMM shows significant difference between baseline and SST IN activation at early but not intermediate ranks (Early: β = -0.147, p = 0.017; Intermediate: β = 0.065, p = 0.090). Right, Fraction of reliable variance explained by descending ranks of PCs for baseline vs. SST IN activation at P14 (right). Significant difference between baseline and SST IN activation at both early and intermediate ranks (Early: β = -0.167, p < 0.001; Intermediate: β = 0.119, p = 0.006).

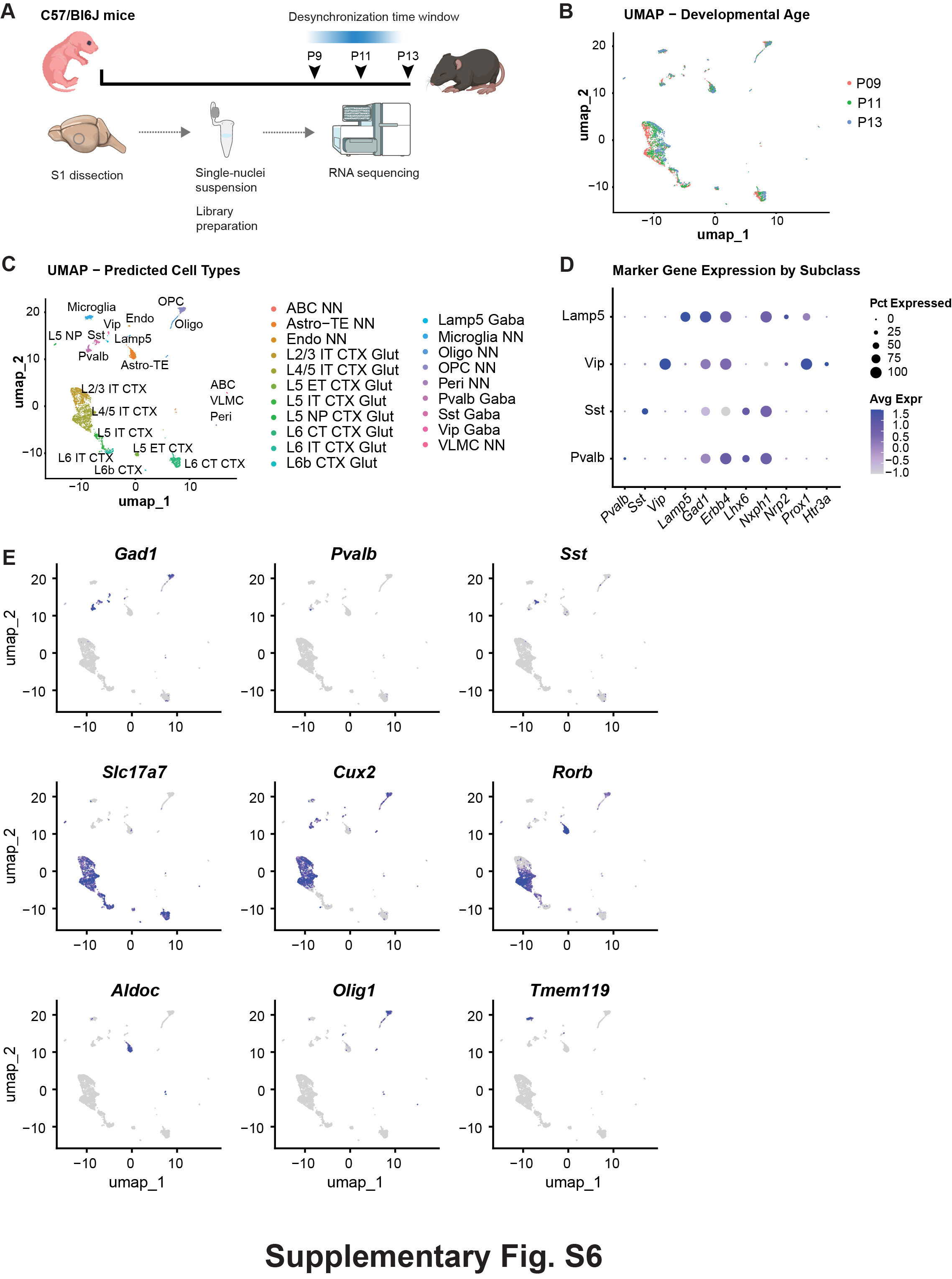

**Supplementary Fig. S6: S1 snRNA-seq dataset validation.**

(A) Schematic for S1 dissection and snRNA-seq (Images from NIH BioArt).

(B) UMAP colored by developmental age.

(C) UMAP colored by predicted cell types.

(D) Dot plot with canonical inhibitory neuron markers.

(E) Feature plots with markers.

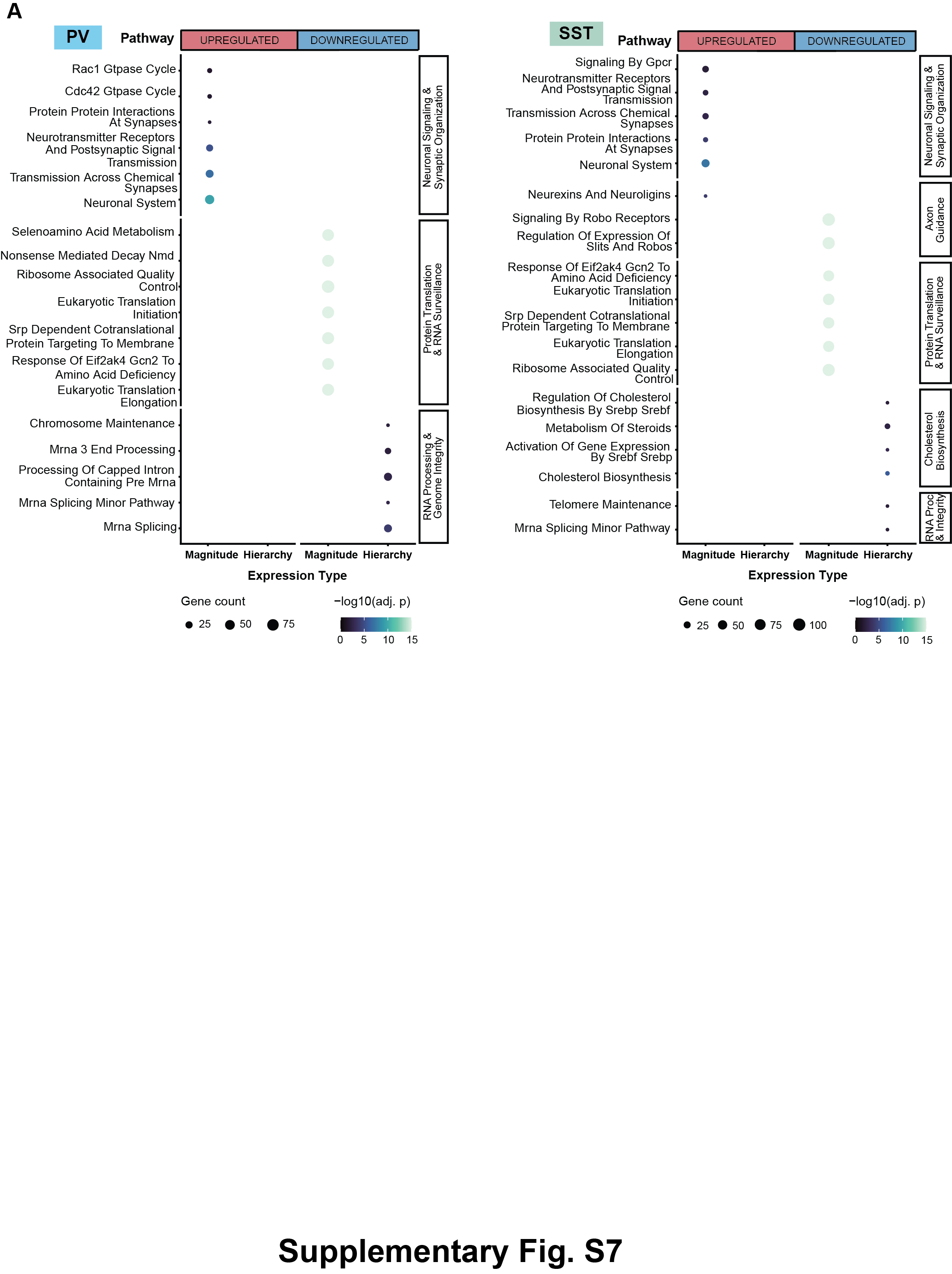

**Supplementary Fig. S7: Pathways upregulated and downregulated in PV and SST INs between P8 and P11.**

(A) Top pathways from over-representation analysis performed using MsigDB Reactome gene sets for PV INs (left) and SST INs (right). Upregulated pathways are associated with changes in expression magnitude and include neuronal signaling and synapse organization pathways while downregulated pathways include both changes in gene expression hierarchy and gene expression magnitude.

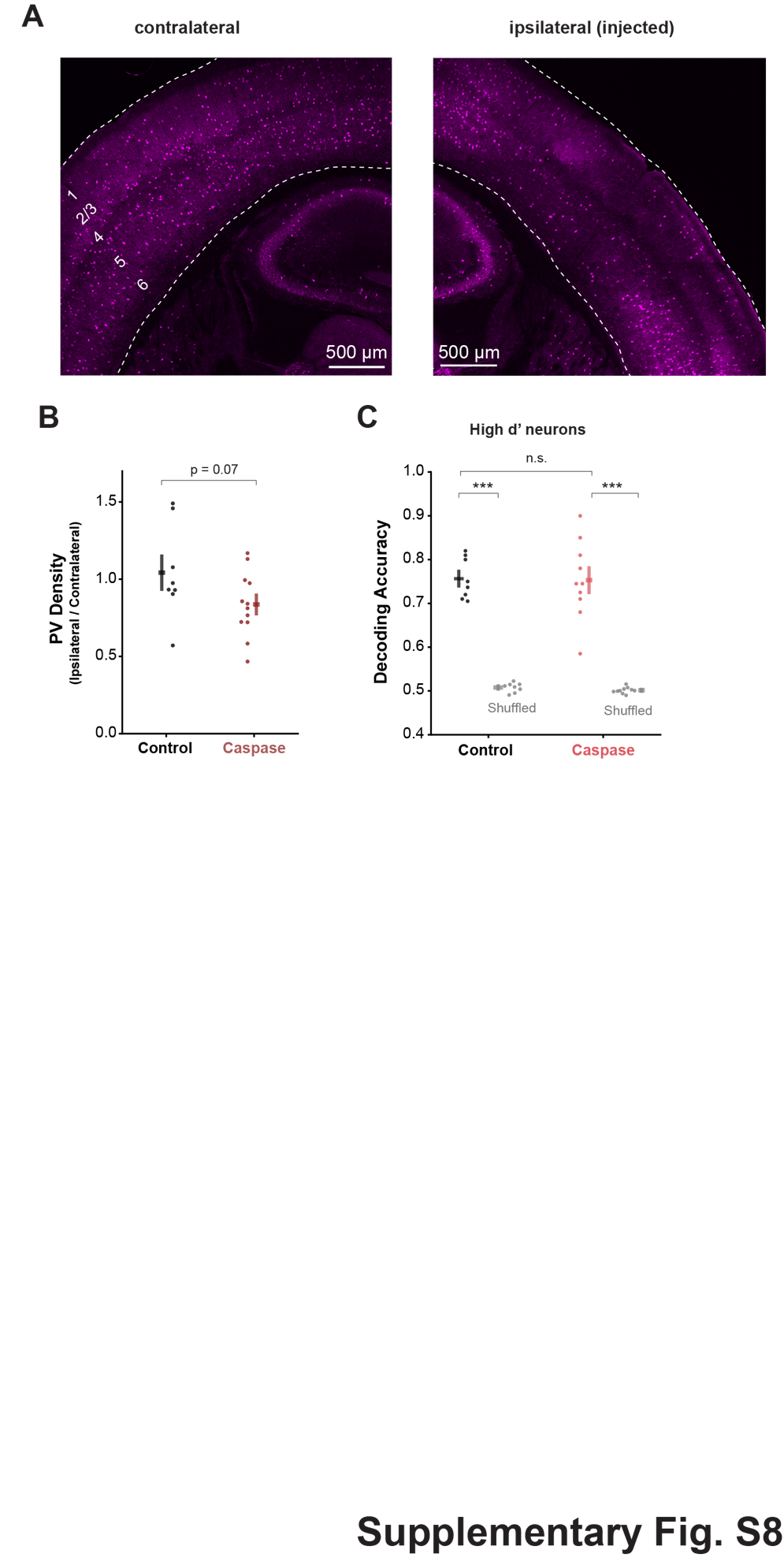

**Supplementary Fig. S8: Viral-mediated caspase-3 reduces PV IN density but does not affect decoding accuracy.**

(A) Immunofluorescent staining for *Pvalb* on the contralateral and ipsilateral hemispheres of S1.

(B) PV IN density decreased in the Caspase-3 injected group (two-sided Mann-Whitney U, p = 0.072).

(C) Decoding accuracy for control and Caspase-3 groups compared to shuffled. LMM showed significant main effects of Shuffling (Wald χ^2^ = 54.65, p = 1.44x10^-13^) but not Group (Wald χ^2^ = 0.40, p = 0.525).

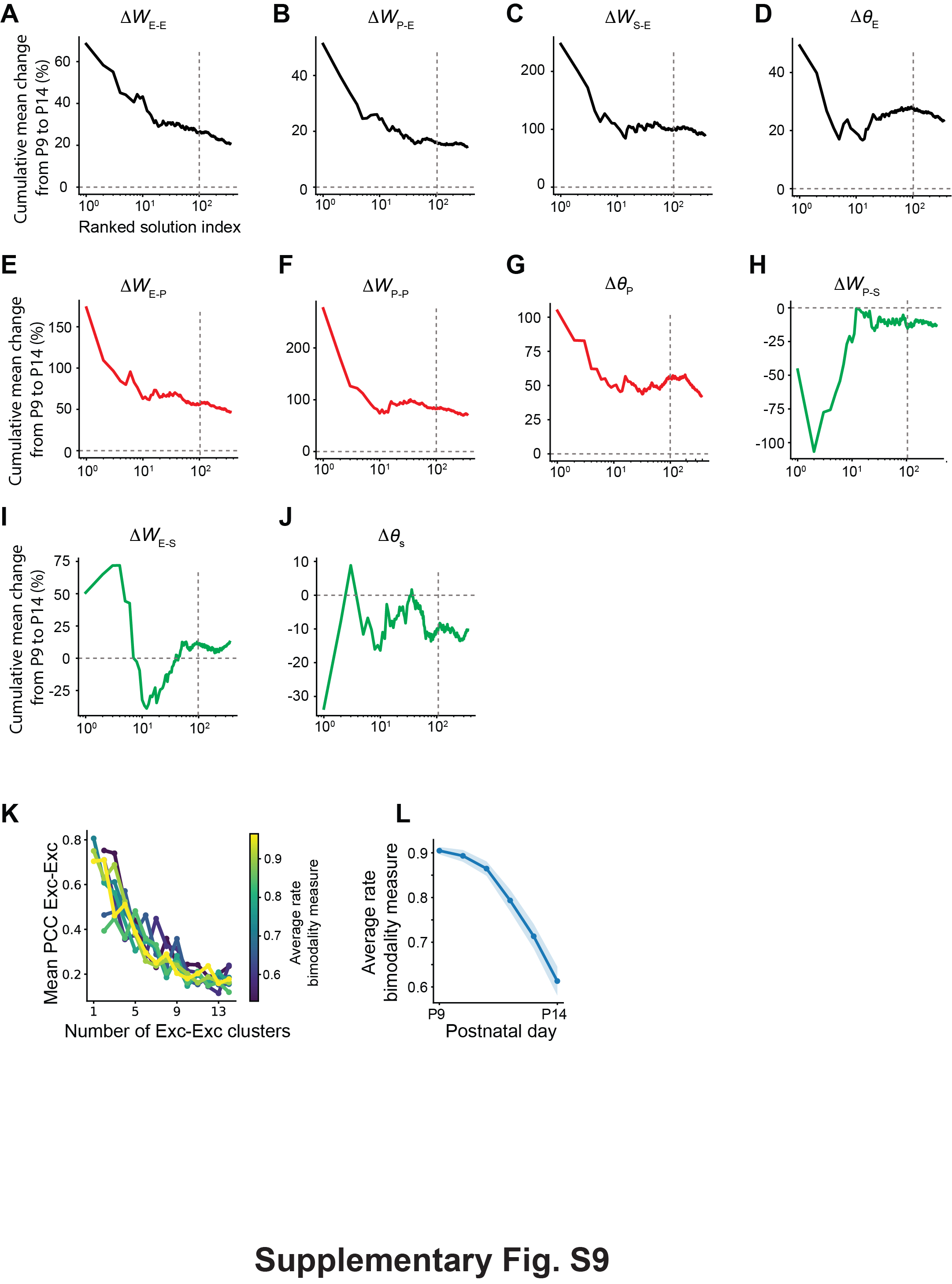

**Supplementary Fig. S9: The parameters’ change is robust across parameter values and the number of clusters impacts the PCC of Excitatory neurons in the SNE dynamical regime, which changes during development.**

(A-D) Synaptic output connections and thresholds: synaptic connections to Exc (A), PV (B), SST (C) and effective firing threshold (D). The vertical dashed line marks the first 100 solutions we considered when evaluating averages over solutions, ranked by biological alignment with development and neural activity statistics for the model parameters’ change from P9 to P14.

(E-H) Same as (A-D), but for PV neurons.

(I and J) Same as (A and D), but for SST. Notice that, for SST neurons, our model does not include projections to PV or SST, in line with biological findings.

(K) The average pairwise PCC between excitatory neurons as function of the number of clusters in the connectivity matrix between excitatory neurons. Each colored line represents a different value of the average bimodality measure of the average firing rate.

(L) Bimodality measure for the average firing rate as a function of development, averaged over the top 100 solutions sorted based on their alignment with developmental parameter change and neural activity statistics.

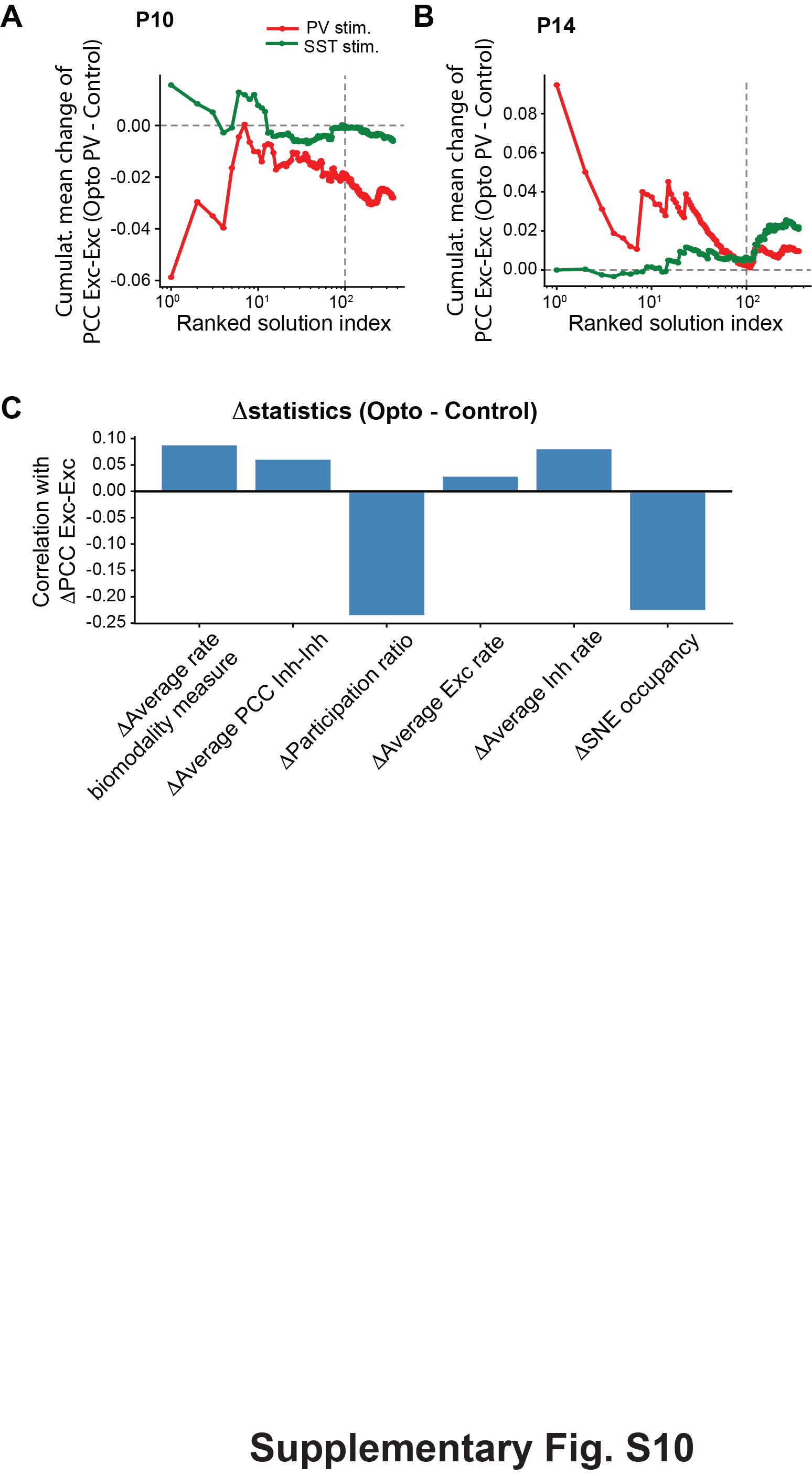

**Supplementary Fig. S10: The predictions of the model are robust across parameter values, and PCC change induced by optogenetic stimulation of PV neurons is mostly correlated with the activity dimension and SNE occupancy.**

(A) Cumulative mean of the change of predicted PCC between control and optogenetics at P10. The dashed black vertical line represents the top 100 solutions, sorted as described in Figure 6C, which are included in the average in Figure 7A and the histograms in Figures 7C and 7D.

(B) Same as A, but for P14.

(C) Correlation between the change in average pairwise PCC among excitatory neurons (baseline - PV stimulation) and the change in several summary statistics of the neural activity as predicted by the model. The SNE occupancy was defined as the product of the SNE mean duration and SNE frequency.
